# HIV and Cocaine exposure promote Tau phosphorylation through RSK-1 in a GSK3β-independent manner

**DOI:** 10.64898/2026.04.14.718541

**Authors:** Adhikarimayum Lakhikumar Sharma, Ilker K. Sariyer, Ulhas P. Naik, Mudit Tyagi

**Author notes:** Correspondence (M.T).

## Abstract

HIV and cocaine are known to disrupt neuronal signaling and contribute to neurocognitive dysfunction, yet the underlying molecular mechanisms are not clear. In this study, we delineate the underlying molecular mechanism by which HIV and/or cocaine enhance Tau phosphorylation (p-Tau S396), a marker of Tau-mediated neuropathies. Furthermore, we elucidate how these two independent neuropathogenic factors, cocaine and HIV, exploit distinct yet convergent signaling pathways to drive this pathological event. We demonstrate that HIV robustly activates and upregulates RSK1, which functions upstream of AKT and promotes Tau phosphorylation through an AKT-independent mechanism while simultaneously inactivating GSK3β via serine-9 phosphorylation (p-GSK3β S9). However, cocaine not only activates RSK1 but also strongly stimulates AKT1, resulting in sustained GSK3β inhibition and persistent Tau phosphorylation. Notably, Tau phosphorylation persists even under conditions of GSK3β inactivation in both HIV and cocaine exposure, revealing a previously unrecognized GSK3β-independent mechanism of Tau modification. Collectively, these findings identify RSK1 as the primary mediator of Tau phosphorylation upon HIV and/or cocaine exposure, and uncover a novel RSK1-driven, GSK3β-independent pathway contributing to Tauopathy. Through a combination of immunofluorescence, immunoblotting, genetic knockout, and overexpression approaches, we establish RSK1 as a central signaling hub linking the AKT-GSK3β pathway to Tau phosphorylation. We demonstrate that RSK1 operates as a critical upstream regulator of AKT and GSK3β signaling, playing dual roles, both activating AKT and suppressing GSK3β, thereby uncovering a novel layer of pathways that regulates Tau phosphorylation. The reproducibility of these main signaling pathways across SH-SY5Y neurons, mixed cell 3D spheroids, and human brain organoids underscores the robustness and biological relevance of this mechanism. Collectively, these findings reveal mechanistic convergence of HIV and cocaine on RSK1-dependent signaling and provide critical insight into how diverse neuropathic / neuropathological factors remodel neuronal signaling to drive Tau-associated dysfunction. These findings provide novel mechanistic insight into the molecular underpinnings of neuro-HIV and substance abuse associated Tauopathy. By identifying RSK1 as a master regulator and demonstrating that Tau phosphorylation can bypass GSK3β inhibition, our study advances understanding of signaling complexity and highlights new opportunities for therapeutic intervention. Targeting RSK1 may represent a promising strategy to mitigate Tau pathology, induced due to insoluble aggregates of phosphorylated Tau, a common factor promoting cognitive decline not only in individuals with Alzheimer’s disease but also in those exposed to cocaine or/and infected with HIV.

**Significances:** This study demonstrates that exposure to HIV and/or cocaine induces Tau phosphorylation at serine 396 (S396), a well-established marker of Tau pathology, and delineates how these two independent neuropathogenic factors engage distinct yet convergent signaling pathways to drive this pathogenic event. We show that HIV exposure drives robust RSK1 activation, positioning it upstream of AKT to promote Tau phosphorylation via an AKT-independent mechanism, while concurrently suppressing GSK3β activity through serine-9 phosphorylation. In contrast, cocaine, while only moderately activating RSK1, primarily enhances AKT signaling, leading to sustained GSK3β inhibition and increased Tau phosphorylation. Notably, Tau phosphorylation persists even under conditions of GSK3β inactivation in both settings, revealing a previously unrecognized, RSK1-centered, GSK3β-independent pathway of Tau modification. Overall, our findings demonstrate that Tau phosphorylation in the context of HIV infection and cocaine exposure is a complex, multi-layered regulatory process involving multiple signaling nodes. Importantly, we identify RSK1 as a central integrative hub linking viral and substance-induced signaling to downstream Tau pathology. This work advances our understanding of the molecular mechanisms underlying neuroHIV and substance abuse–associated neurodegeneration. Furthermore, it highlights RSK1 as a novel and promising therapeutic target for mitigating Tauopathy in both cocaine-using and non-using people with HIV (PWH).

**Highlighted points:**

- RSK1 acts as a central regulator of Tau phosphorylation, capable of driving this process through a GSK3β-independent mechanism.
- HIV promotes Tau phosphorylation primarily via robust upregulation and activation of RSK1, operating largely independent of AKT1, while concurrently inducing GSK3β inactivation.
- Drugs of abuse, such as cocaine induces Tau phosphorylation through dual activation of AKT1 and RSK1, alongside sustained inactivation of GSK3β.
- Tau phosphorylation persists despite GSK3β inhibition, revealing a complex AKT1-RSK1 signaling axis and underscoring the dominant role of GSK3β-independent mechanisms in Tau pathology following HIV and cocaine exposure.
- HIV and cocaine engage distinct yet convergent signaling pathways that disrupt neuronal homeostasis and drive tauopathy, providing mechanistic insight into neuroHIV and substance abuse-associated neurodegeneration.
- RSK1 functions as a key upstream modulator of AKT and GSK3β pathways, positively regulating AKT signaling while negatively regulating GSK3β activity.
- RSK1 emerges as a potential therapeutic target, offering new opportunities for intervention in HIV-associated neurocognitive disorders (HAND) and drug-induced neurodegeneration.
- Established and characterized H80 cells as a novel neuronal cell model and demonstrated their suitability for studying neuron-specific signaling pathways, including Tau phosphorylation.
- The conserved and widespread nature of the signaling cascade driving Tau phosphorylation in response to HIV and/or cocaine exposure was validated across multiple model systems, including both 2D neuronal cell cultures and 3D systems such as human brain organoids and spheroids.

**Strength of the Study:** This original study provides novel mechanistic insight into how HIV and cocaine, two independent neuropathological factors, converge and diverge on intracellular signaling pathways to regulate Tau phosphorylation. By integrating immunofluorescence, immunoblotting, genetic knockout, and overexpression approaches, we identified RSK1 as a master regulator of Tau phosphorylation. Importantly, we discovered that HIV robustly upregulates and activates RSK1 to promote Tau phosphorylation through an AKT-independent route while simultaneously inactivating GSK3β. On the other hand, cocaine exerts a moderate effect on RSK1 but strongly stimulates AKT to induce GSK3β inactivation and drive Tau phosphorylation. A key strength of this work is the discovery that Tau phosphorylation persists despite GSK3β inactivation, revealing a complex, GSK3β-independent mechanism, involving RSK1 in Tau pathology. Moreover, our study, for the first time, identify RSK1 as an upstream regulator of AKT-GSK3β signaling cascade, enhancing AKT signaling while simultaneously inhibiting GSK3β activity, thereby underscoring the critical role of RSK1 in Tau phosphorylation and associated illnesses, such as HAND and Alzheimer’s disease. Together, these findings not only advance our understanding of the molecular underpinnings of neuroHIV and substance abuse associated tauopathy but also highlight RSK1 as a promising therapeutic target for not only HIV and cocaine induced neurotoxicity but also other neurodegenerative diseases, such as Alzheimer’s disease. Another key strength of this study is the establishment and characterization of H80 cells as a novel neuronal model, demonstrating their suitability for investigating neuron-specific signaling pathways, including Tau phosphorylation. The combination of comparative signaling analysis, genetic perturbations, and integrative mechanistic modeling makes this study both conceptually and technically novel, besides broadly relevant to the fields of neurovirology, addiction neuroscience, neurodegeneration, and cognitive impairments.

## Introduction

Human immunodeficiency virus (HIV) infection remains a significant global health concern, with an estimated 38 million people currently living with the virus worldwide [1]. Although the introduction of combination antiretroviral therapy (ART) has markedly improved life expectancy and viral suppression in people with HIV (PWH), the burden of HIV-associated neurocognitive disorders (HAND) persists [2, 3]. HAND affects approximately 30-50% of PWH, even in those achieving robust viral suppression on ART [4]. The etiology of HAND is multifactorial and complex, involving persistent neuroinflammation, the activity of neurotoxic viral proteins (e.g., Tat, gp120), and dysregulation of host signaling pathways that collectively disrupt synaptic integrity and neuronal function [3]. Importantly, while neurons are the primary cells affected in HAND, glial cell population in the central nervous system (CNS), are increasingly recognized as key mediators of HAND-related neuropathology [5, 6].

The microtubule-associated protein Tau is a central regulator of neuronal function, ensuring the stability of axonal microtubules and supporting efficient transport of cargo essential for synaptic activity and neuronal survival [7, 8]. Under physiological conditions, Tau protein undergoes tightly regulated cycles of phosphorylation and dephosphorylation that allow dynamic modulation of cytoskeletal structure. However, disruption of this phosphorylation event (Tau hyperphosphorylation) gives rise to neurodegenerative disease [7, 9, 10]. Aberrantly phosphorylated Tau exhibits diminished binding to microtubules, misfolds into abnormal conformations, and progressively accumulate into insoluble neurofibrillary tangles [11, 12]. These inclusions not only serve as histopathological hallmarks of Alzheimer’s disease (AD) and related tauopathies but also correlate strongly with synaptic dysfunction, neuronal loss, and the severity of cognitive decline [13, 14]. Multiple factors contribute to the pathological transformation of Tau, ultimately driving its involvement in AD and related dementias [15, 16]. Emerging evidence suggests that HIV infection, even in individuals receiving suppressive antiretroviral therapy, can disrupt the physiological regulation of Tau phosphorylation. Since neurons do not express canonical HIV entry receptors such as CD4 and co-receptors CCR5 or CXCR4, they are not infected by the virus [17, 18]. Nevertheless, neuron cells remain profoundly susceptible to the downstream consequences of viral exposure. Instead of direct infection, neuronal injury arises predominantly through indirect mechanisms, most notably the actions of soluble viral proteins, mainly Tat and gp120, and released cytokines from infected glial or immune cells [19, 20]. These cytotoxic factors dysrupt cellular homeostasis and interfere with host kinase-phosphatase signaling cascades, leading to dysregulated Tau phosphorylation events that compromise cytoskeletal integrity, axonal transport, and synaptic stability ultimately leading to HAND, and related tauopathies.

Cocaine, one of the most prevalent substances abused among PWH, is a well-established cofactor in the progression of HAND [21, 22]. Cocaine abuse independently exacerbates neurodegenerative processes, accelerates cognitive decline, and has been associated with increased susceptibility to HIV infection and replication within the CNS [22–24]. Mechanistic studies suggest that cocaine induces oxidative stress, disrupts blood-brain barrier integrity, activates glial cells, and modulates multiple signaling pathways, including those governing inflammation and cell survival. Our recent studies further reveal that cocaine use enhances HIV transcription by activating transcription factors such as NF-κB and MSK1, altering epigenetic modifications at the long terminal HIV repeat (LTR) promoter [24, 25].

Beyond transcriptional activation, we have shown that cocaine increases the susceptibility of CD4⁺ T cells to HIV infection by augmenting key co-stimulatory signaling pathways, involving, NF-kB, NFAT and AP-1, thereby creating a cellular state more favorable to viral entry and replication [25–27]. Additionally, we have demonstrated that cocaine activates DNA-dependent protein kinase (DNA-PK) in both T cells and microglial cells, which alleviates RNA polymerase II pausing at the LTR. This effect is mediated through phosphorylation of TRIM28, a chromatin-associated repressor, thus enabling more efficient transcriptional elongation and sustained viral gene expression [28, 29]. These molecular changes establish a favorable environment for persistent HIV activity and may synergize with host signaling dysregulation to exacerbate neuropathology, particularly within the central nervous system.

Neurodegeneration is the consequence of dysregulated intracellular signaling, in which kinases play a crucial role [30]. In the healthy normal brain, a balance between kinases and phosphatases ensures proper regulation of cytoskeletal dynamics, synaptic activity, and stress adaptation. Disruption can cause series of phosphorylation events that promote neuronal dysfunction [31]. Therefore, several cellular pathways have been involved in the regulation of Tau phosphorylation ultimately leading to neurodegeneration. However, glycogen synthase kinase 3 beta (GSK3β), a serine/threonine kinase that directly phosphorylates Tau at multiple pathological sites remain a major kinase [32, 33]. GSK3β activity is inhibited by phosphorylation at serine 9 (Ser9), a modification typically mediated by the upstream kinase AKT (also known as protein kinase B), a central node in cell survival, metabolism, and growth signaling [34]. Dysregulation of this AKT-GSK3β axis has been consistently reported in models of tauopathy, and other neurodegenerative conditions, underscoring its pathogenic significance [35, 36]. In addition to GSK3β, several other kinases have been shown to phosphorylate Tau, including Cyclin-dependent kinase 5 (CDK5), Extracellular signal-regulated kinases (ERK1/2), c-Jun N-terminal kinase (JNK), p38 MAPK, Microtubule affinity-regulating kinases (MARKs), AMP-activated protein kinase (AMPK) [37, 38], Protein kinase A (PKA), Calcium/calmodulin-dependent kinase II (CaMKII) [39], and Protein kinase C (PKC), act on overlapping sets of phosphorylation sites and often respond to cellular stress signals such as inflammation, oxidative damage, and excitotoxicity. Dysregulation of the MAPK/ERK signaling pathway has been strongly associated with neurodegenerative disorders, including AD, where abnormal kinase activity contributes to synaptic dysfunction, Tau hyperphosphorylation, and neuronal loss [40, 41].

Ribosomal S6 kinase 1 (RSK1, encoded by *RPS6KA1*) is known to be a key downstream effector of MAPK/ERK pathway and plays important roles in regulating cell growth, survival, and gene expression [42, 43]. Despite its central role in MAPK/ERK signaling, RSK1 has not yet been systematically investigated in the context of Tau phosphorylation or AD, and direct evidence linking its dysregulation to disease onset or progression remains limited. This gap in knowledge prompted us to investigate this aspect in detail and define if RSK1 is an underexplored contributor to the molecular mechanisms underlying Tauopathy responsible for AD and related neurodegenerative conditions. Additionally, several of the aforementioned kinases have been extensively studied in AD and other tauopathies, their specific contributions to Tau dysregulation during HAND remain poorly understood. Viral proteins such as Tat and gp120 are known to disrupt intracellular signaling, yet the precise mechanisms by which they interact with the kinase-phosphatase networks that regulate Tau remain unclear. Elucidating the signaling pathways through which Tau pathology accelerates HAND is therefore a critical research priority. Addressing this knowledge gap will not only advance our mechanistic understanding of HAND pathogenesis but may also reveal convergent therapeutic targets relevant to both classical and virally mediated neurodegenerative disorders, including AD and HAND.

Selecting an appropriate model system is critical for studying neurodegenerative diseases, as it directly influences the reliability, reproducibility, and translational relevance of the findings. While primary neurons offer high physiological relevance, they are short-lived, fragile, and technically challenging to maintain under *in vitro* conditions, highlighting the need for alternative neuron-like systems that are more experimentally tractable yet retain key neuronal features [44]. Glioma cell lines provide a robust, practical, and experimentally tractable platform to investigate mechanisms of neurodegeneration due to their neural origin, robust growth properties, and retention of signaling pathways relevant to function and disease pathology [45, 46]. Unlike primary neurons, which are post-mitotic and difficult to maintain long term, glioma cells readily expand *in vitro*, enabling reproducible experiments and large-scale molecular and pharmacological studies [47]. Importantly, these cells retain critical signaling pathways relevant to disease pathology, such as MAPK/ERK and PI3K/AKT signaling, oxidative stress responses, and glial-neuronal interactions, all of which are central to the progression of neurodegenerative disorders [48–50]. Since glial dysfunction and altered kinase signaling contribute significantly to synaptic loss, protein aggregation, and neuronal death, glioma cells serve as a practical surrogate model to dissect these mechanisms. Although glioma cells cannot fully replicate the complexity of the CNS, especially neuronal and glial interactions *in vivo*, their tractability and physiological relevance make them a useful and valuable model for mechanistic studies of neurodegeneration, as well as testing the potential therapeutics.

In this study, using both 2D and 3D neuronal model systems, we delineate the molecular mechanisms underlying Tau phosphorylation in response to HIV infection and/or cocaine exposure. We demonstrate that HIV exposure robustly upregulates and activates RSK1, which in turn inactivates GSK3β through an AKT-independent mechanism. Activated RSK1 directly promotes Tau phosphorylation. On the other hand, cocaine exposure not only induces RSK1 but also strongly activates AKT1, leading to GSK3β inactivation through phosphorylation at serine 9 (p-GSK3β S9) in an AKT-dependent manner, thereby further enhancing Tau phosphorylation. Notably, Tau phosphorylation persists even under conditions of GSK3β inhibition during both HIV and cocaine exposure, indicating that Tau modification is primarily driven through an RSK1-centered, GSK3β-independent pathway. These findings highlight the pivotal role of RSK1 and underscore the complexity of the signaling networks regulating Tau phosphorylation. Using complementary genetic and pharmacological approaches, including CRISPR/Cas9-mediated knockout, overexpression systems, and selective kinase inhibition, we further establish that RSK1 functions upstream of both AKT activation and GSK3β inactivation, exerting context-dependent effects on Tau phosphorylation. Collectively, our findings highlight the kinase signaling crosstalk underlying HAND and cocaine-associated tauopathy, identifying RSK1 as a mechanistic hub and potential therapeutic target for neurotoxicity and HAND in PWH, including those who use illicit substances, such as cocaine.

## Materials and Methods

### Cell Culture

H80 cells (originally obtained from the Darell Bigner Laboratory, Duke University) [51] were maintained in DMEM/F12 medium supplemented with 10% fetal bovine serum (FBS) and 1% penicillin–streptomycin. Jurkat T cells (human CD4⁺ T lymphocyte line; ATCC TIB–152), MT-4 cells and U937 cells were cultured in RPMI–1640 supplemented with 10% FBS, 1% penicillin–streptomycin, and 2 mM L–glutamine. HEK293T cells, microglial cells, and SH–SY5Y neuroblastoma cells were propagated in Dulbecco’s modified Eagle medium (DMEM) containing 10% FBS, 1% penicillin–streptomycin, and 2 mM L–glutamine. All cell lines were maintained at 37 °C in a humidified incubator with 5% CO_2_ and were used between passages 3 and 8.

### Inhibitor Treatments

H80 cells [51] were seeded and allowed to adhere overnight prior to treatment. Cells were incubated with selective inhibitors targeting RSK1 or GSK3β, at a final concentration of 10 µM each. The RSK1 inhibitor BI-D1870 (Selleckchem, Cat. No. S2843), and GSK3β inhibitor CHIR99021 (Tocris Bioscience, Cat. No. 4423), were prepared as stock solutions in Dimethyl sulfoxide (DMSO) and prepared working solutions at 10 mM. Cells were treated with inhibitors or equivalent volumes of DMSO vehicle control for 24 hours at 37°C in a humidified 5% CO₂ incubator. Following treatment, cells were either exposed to HIV or cocaine or both and harvested for downstream analyses including Immunoblotting. All treatments were performed in experimental triplicate or biological triplicate to ensure reproducibility.

### Cocaine Treatment

Cells were treated with 10 µM cocaine hydrochloride. For acute exposure, treatments were applied for durations ranging from 15 minutes up to 6 hours. Otherwise, specifically mentioned all the treatments are done chronically. For chronic exposure, cells received two treatments each day randomly for 48 hours and at least 30 min-3h prior cells harvesting. Control cells were treated with PBS or kept untreated.

### HIV Virus Production and Infection

Jurkat cells or MT-4 cells were infected with replication-competent Human Immunodeficiency Virus Type 1 (strain 93/TH/051, R5- and X4-tropic virus) (NIH AIDS Reagent Program) by spinoculation at 1,200 × g for 2 h at 25°C in the presence of 8 µg/mL Polybrene. Following infection, cells were incubated for 48 h-72 h. Supernatants containing HIV virions were harvested, cleared by low-speed centrifugation (500 × g, 10 min), filtered through a 0.45 µm syringe filter, and stored at −80 °C. HIV production was confirmed by immunoblotting for the HIV p24 capsid protein.

### H80 Exposure to HIV

H80 were seeded and exposed to HIV-containing supernatant or in normal medium (control) by spinfection at 1,000 rpm for 2 h at room temperature (RT). The following day, cells underwent a second spinfection under the same conditions and were subsequently transferred to 100-mm dishes. After 48 h, cells were harvested for protein analysis. Control H80 cells were processed identically with supernatant from uninfected Jurkat/MT-4 cells.

### Lentiviral production and CRISPR/Cas9-mediated RPS6KA1 knockout

Lentiviral particles encoding Cas9 were produced by co–transfecting HEK293T cells with either lentiCRISPR v2 (Addgene #52961) or lentiCas9–Blast (Addgene #52962), a gift from Feng Zhang [52], together with the packaging plasmid psPAX2 (Addgene #12260), and the envelope plasmid pMD2.G (Addgene #12259), a gift from Didier Trono using Lipofectamine 2000 (Thermo Fisher Scientific). Viral supernatants were harvested 48 h post–transfection, clarified through a 0.45–µm filter, and used immediately or stored at −80 °C. H80 cells were transduced in the presence of 8 µg/mL polybrene and selected with puromycin (1–2 µg/mL)/ blasticidin to generate stable Cas9–expressing populations. To disrupt RPS6KA1, Cas9–positive H80 cells were subsequently transduced with lentiviral particles encoding one of three independent RPS6KA1–targeting sgRNAs (BRDN0001148481, Addgene #75499; BRDN0001145974, Addgene #75497; BRDN0001148103, Addgene #75498), originally developed by John Doench and David Root [53]. These sgRNA lentiviruses were produced in HEK293T cells using the same Lipofectamine–based system. Following transduction, cells were allowed to recover for 48 h and then selected with puromycin (1–2 µg/mL) for 3–5 days to obtain RSK1–knockout populations.

### Overexpression in H80 Cells

For RSK1 overexpression, H80 cells were seeded at 60–70% confluence in 60 mm plates and transiently transfected with 2 µg of a CMV promoter-driven full-length human RSK1 expression plasmid (pKH3-human RSK1, Addgene cat no #13841, a gift from John Blenis [54]) or empty vector control using Lipofectamine 2000 (Thermo Fisher Scientific) according to the manufacturer’s instructions. Briefly, plasmid DNA and Lipofectamine reagent were diluted separately in Opti-MEM (Gibco), combined, and incubated for 30 minutes at RT before adding drop by drop to cells. Cells were incubated with the transfection complexes for 6 hours, after which the medium was replaced with fresh growth medium. Protein lysates were harvested 48 hours post-transfection using lysis buffer with protease and phosphatase inhibitors (Roche). Overexpression efficiency was confirmed by immunoblotting with anti-RSK1 antibodies.

### Spheroids formation

Three-dimensional spheroids were generated using 96-well round-bottom Biofloat 3D cell culture plates (Sarstedt, Cat. No. 83.3925.400), which provides a non-adhesive surface to promote uniform spheroid formation. To prevent cell attachment, each well was pre-treated with 60 µL of Anti-Adherence Rinsing Solution (AARS; Stemcell Technologies, Cat no #07010) and incubated under sterile conditions at RT for 24 h. Following incubation, the AARS was aspirated and stored it for potential reuse, and wells were rinsed with 100 µL of phosphate-buffered saline (PBS) to remove residual solution. Prepared plates were either used immediately or stored in sterile bags at 4 °C for up to two weeks. For spheroid assembly, a mixed cell suspension containing H80 cells, SH-SY5Y neuroblastoma cells, and microglia (5,000 cells of each type) was prepared in 150 µL of DMEM supplemented with 10% FBS and 1% penicillin–streptomycin. This suspension was dispensed into each treated well, and plates were incubated at 37 °C in a humidified atmosphere with 5% CO₂ for 24 h to allow initial aggregation and spheroid formation. After 48 h of culture, 100 µL of medium was carefully removed from each well and replaced with fresh medium containing HIV virus to initiate infection or exposure. Spheroids were incubated for 5 h under the same conditions, after which the medium was exchanged for fresh medium containing either cocaine or no treatment. Cocaine was administered twice daily for 48 h to mimic repeated exposure. At the end of the treatment period, spheroids were harvested by pooling 24 spheroids into a single Falcon tube, representing one biological sample. A total of 96 spheroids were collected, corresponding to four experimental groups: untreated control (24 spheroids), cocaine-treated (24 spheroids), HIV-exposed/infected (24 spheroids), and HIV-exposed/infected plus cocaine-treated (24 spheroids) (spheroid figure in Supplementary). Each pooled sample was washed with 1 mL PBS to remove residual medium and treatment compounds, followed by addition of 80 µL of 1× passive lysis buffer (Promega E1941) to facilitate cell lysis and protein extraction for Immunoblot analyses.

### Generation of human cerebral organoids (hCOs)

Human cerebral organoids (hCOs) were generated from human induced pluripotent stem cells (hiPSCs) following our previously established protocols, as described in detail in reference [55]. Briefly, human induced pluripotent stem cells (hiPSCs), derived from dermal fibroblasts, were used to generate hCOs following STEMdiff™ protocols (STEMCELL Technologies). The cells were plated in ultra-low attachment 96-well plates at 11,000 cells per well and incubated for 24 hours to form embryoid bodies (EBs). As the EBs grew to 400–600 μm over about 5 days, they were transferred to 24-well plates and cultured in induction medium for 48 hours. The EBs were then embedded in Matrigel and moved to 6-well plates with expansion medium, where they developed neuroepithelial structures after 3 days. Finally, the organoids were matured on an orbital shaker at 70 rpm in maturation medium at 37°C for an additional 40 days.

### Immunoblotting

Total cell lysates were prepared using 1X Passive Lysis Buffer (Promega E1941) supplemented with protease and phosphatase inhibitor cocktails (Roche), following the manufacturers’ instructions. Following cell harvesting, lysates were incubated on ice for 30 minutes with intermittent vortexing for 30 seconds every 10 minutes to facilitate complete lysis. For the Spheroids and organoids, samples were lysed by mechanical disruption with passive lysis buffer through repeated passage through a 200 µL pipette tip, followed by eight cycles of rapid freeze–thaw in liquid nitrogen and a 37 °C water bath. The lysates were then incubated on ice. After incubation, samples were centrifuged at maximum speed (≥14,000 × g) for 30 minutes at 4 °C to pellet cell debris. The resulting supernatants were collected, and protein concentrations were determined using the Pierce™ BCA Protein Assay Kit (Thermo Fisher Scientific). Protein concentration was normalized, and an equal amount of protein was mixed with 5X Laemmle Sample buffer, heated to 95°C for 10 min, and then resolved by SDS-PAGE on a 9% or 10% or 12% gel at 120 volts until the dye reached the bottom. The resolved proteins were then transferred to a nitrocellulose membrane (Amersham). Membranes were blocked for 1 h at RT in 3% bovine serum albumin (BSA) in Tris-buffered saline containing 0.1% Tween-20 (TBS-T), followed by overnight incubation at 4 °C with primary antibodies against phospho-RSK1 Ser380 (sc-136476), phospho-RSK1 thr348 (sc-101770), phospho-p90RSK (Thr359/Ser363) (CST#9344), RSK1 (CST #9347), RSK1/2/3 (CST #14813), phospho-AKT T308 (CST #4056), phospho-AKT S473 (CST #4060), AKT1 (CST #2938), phospho-GSK3β S9 (CST #5558), GSK3β (CST #12456), phospho-Tau (CST #9632S), Tau (CST #46687), MAP2 (17490-1-AP), GAPDH (sc-25778), and β-actin (Sigma-Aldrich A5316). After three washes with 1X TBST, the blot was detected using the Odyssey infrared imaging system application software 3.0 (Li-cor Bioscience).

### RNA Extraction and Quantitative PCR (qPCR)

Total RNA was extracted from H80 cells after 24 h of HIV exposure using the RNeasy Plus Mini Kit (Qiagen) following the manufacturer’s protocol, ensuring elimination of genomic DNA contamination. RNA purity and concentration were confirmed by nanodrop and RNA gel electrophoresis. Complementary DNA (cDNA) was synthesized from 1 µg of total RNA using the High-Capacity cDNA Reverse Transcription Kit (Applied Biosystems). Quantitative PCR was performed using SYBR Green on a QuantStudio 5 Real-Time PCR system (Applied Biosystems) with gene-specific primers for IL-1β, TNF-α, RSK1, and GAPDH as an internal control. Relative gene expression was quantified by the 2^−ΔΔCt method, normalizing target gene expression to GAPDH and comparing HIV exposed to controls (exposed without HIV). All reactions were conducted in technical triplicates across at least three biological replicates.

### Immunofluorescence staining and imaging

To characterize the H80 cells, H80 cells were cultured on sterile coverslips, which was initially treated with PolyD Lysine and allowed to adhere overnight. Cells were fixed with 4% paraformaldehyde (PFA) in PBS for 30 minutes at RT, followed by permeabilization with 0.25% Triton X-100 in PBS for 10 minutes at RT. After permeabilization, cells were washed thrice with PBS and then incubated with a blocking solution containing 10% horse serum and 2% BSA in PBS for 60 minutes at RT to reduce non-specific binding. Subsequently, cells were washed and then incubated overnight at 4°C with directly conjugated primary antibodies: anti-NeuN Alexa Fluor 647 (Cat. No. 608453, BioLegend), p-Tau S396 (Phospho-Tau (Ser396) (PHF13) Mouse mAb #9632)/anti-Tau phospho ser396 (BioLegend #829001) and MAP2 Polyclonal antibody (proteintech, 17490-1-AP). The following day, samples were washed three times with PBS, incubated with secondary for 1 h and counterstained with Hoechst (300 nM in PBS) for 10 minutes at RT, followed by an additional three PBS washes. Coverslips were mounted using Aqua-Mount mounting medium (Epredia, Cat. No. 13800) and imaged using an EVOS M7000 Imaging System (Cat no. AMF7000) equipped with 20× and 40× oil immersion objectives.

To investigate the cellular effects of cocaine and HIV exposure, exposed cells were fixed with 4% PFA in PBS for 15 minutes at RT. After washing the cell thrice in PBS, Fixed cells were permeabilized with 0.25% Triton X-100 in PBS for 10 minutes at RT, then incubated for 1 hour in blocking buffer composed of 10% horse serum and 2% BSA in PBS. Primary antibodies against phosphorylated Tau (Phospho-Tau (Ser396) (PHF13) Mouse mAb #9632), RSK-1 (CST #9333), p-GSK3β S9 (CST #5558), p-AKT S473 (CST #4060), AKT1 (CST #2938), were incubated overnight at 4°C. The next day, cells were washed thoroughly to remove unbound antibodies and subsequently incubated with species-specific, fluorophore-conjugated secondary antibodies for 45 minutes at RT in the dark. Nuclei were counterstained with Hoechst for 10 minutes at RT, followed by three PBS washes. Coverslips were mounted using Aqua Mount (Epredia, cat. no. 13800), and samples were imaged using an EVOS imaging system equipped with 10x, 20x and 40x objectives.

### Flowcytometry

H80 cells were analyzed for surface expression of the HIV entry receptors CD4, CCR5, and CXCR4 using multicolor flow cytometry. Cells were harvested, washed with PBS containing 2% FBS, and incubated with fluorochrome–conjugated monoclonal antibodies for 30 minutes at 4 °C in the dark. To assess co–expression of CD4 and CXCR4, cells were stained with APC anti–human CD4 (BioLegend, cat. no. 317416) and PE anti–human CD184 (CXCR4) (BioLegend, cat. no. 306505). For CD4 and CCR5 co–staining, cells were incubated with PE anti–human CD4 (BioLegend, cat. no. 357403) and APC/Cyanine7 anti–human CD195 (CCR5) (BioLegend, cat. no. 359110). Following staining, cells were washed, resuspended in PBS, and analyzed on a flow cytometer. Data acquisition and compensation were performed using standard instrument settings, and analysis was conducted with FlowJo software (BD Biosciences).

### Densitometry and Statistical Analysis

All experiments were performed with a minimum of three independent biological replicates and/or experimental triplicates. Immunoblots were quantified using ImageJ (NIH, Version 1.53e). Band intensities were normalized to β-actin or GAPDH or corresponding total protein and expressed as fold change relative to controls. Data are shown as mean ± standard deviation (SD) from ≥3 independent experiments. Statistical analyses were performed using GraphPad Prism v9 (Version 9.1.2). For comparisons between two groups (e.g., control vs. RSK1 knockout or Ctrl vs. RSK1O/E), unpaired two-tailed Student’s T-tests were employed. For experiments involving multiple conditions or time points, one-way or two-way analysis of variance (ANOVA) followed by Dunnett’s multiple comparisons test was used to assess significance, with p < 0.05 considered statistically significant.

## Results

### H80 Cells Exhibit Neuronal Characteristics as Evidenced by NeuN, MAP2, and Tau Expression

Given the considerable variability, as well as the growth and maintenance challenges associated with commonly used neuronal cell lines such as SH-SY5Y [56], we sought to evaluate whether a glioma cell line, H80 retains key neuronal characteristics, particularly relevant signaling pathways and susceptibility to neurotoxicity. Our findings indicate that H80 cells exhibit features suitable for modeling neuron-specific signaling events, including pathways involved in Tau phosphorylation. Importantly, H80 cells offer several practical advantages over conventional neuronal models, including robust and reproducible growth, rapid proliferation, low baseline cytotoxicity, and stable culture behavior, making them a reliable and efficient system for mechanistic studies of neuronal signaling and neurotoxicity. The neuronal characteristics of the H80 glioma cells [51] were confirmed by performing immunofluorescence staining using NeuN, a nuclear marker widely recognized for its specificity to post-mitotic neurons. Both unstained and secondary-only controls were included in parallel to validate antibody specificity and to exclude background artifacts. Our analysis revealed robust nuclear NeuN immunoreactivity across the majority of H80 cells, providing clear evidence that H80 is a neuronal cell line (**Figure 1A**). The expression of Tau, a microtubule-associated protein that plays a critical role in axonal stability and is centrally implicated in tauopathies and other neurodegenerative processes, further substantiates the neuronal identity of H80 cells (**Supplementary Figure S1**). The presence of Tau not only reinforces the neuronal-like characteristics of H80 cells but also highlights their relevance as a model system for studying Tau-associated signaling pathways and neurotoxicity.

**Figure 1:**
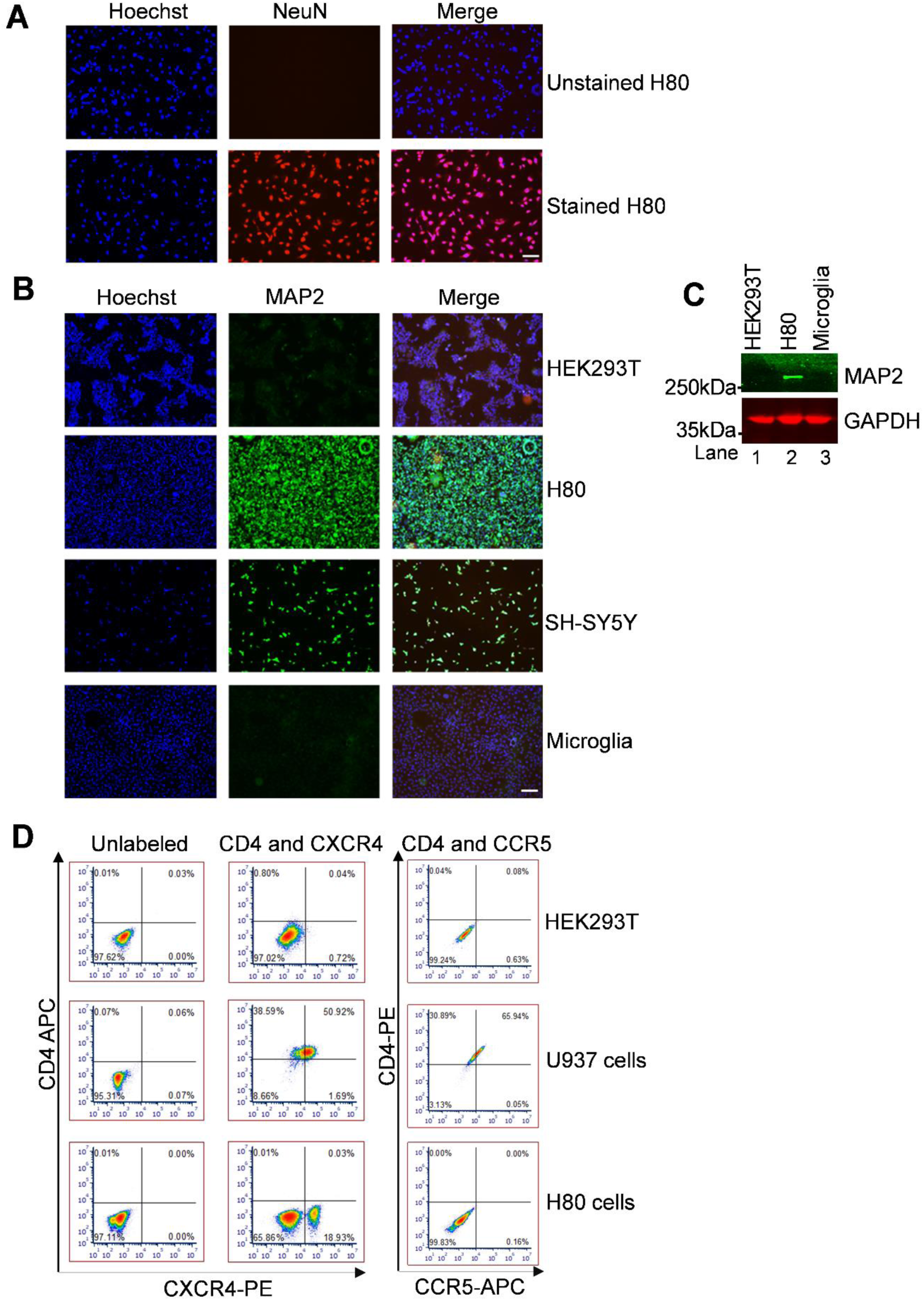
Neuronal characteristics and HIV co-receptor profile of H80 cells. **(A)** Immunofluorescence staining of H80 cells for NeuN using specific primary antibodies, with Hoechst (blue) counterstaining and unstained as controls. NeuN was detected, confirming the neuronal-like phenotype of H80 cells. **(B)** Comparative immunofluorescence staining of H80 cells, microglial cells, SH-SY5Y cells (positive control), and HEK293T cells (negative control) demonstrated that MAP2 expression was restricted to H80 and SH-SY5Y cells. The inclusion of HEK293T as a negative control and SH-SY5Y as a positive control confirmed assay specificity. Detection of MAP2 in H80 cells indicates that these cells exhibit features of differentiated neurons. **(C)** Immunoblotting verified MAP2 expression in H80 cells but not in HEK293T or microglial cells, further supporting the neuronal identity of H80 cells. Collectively, these findings demonstrate that H80 cells express canonical neuronal markers, including NeuN and MAP2, positioning them as a relevant model for investigating neuron-related molecular mechanisms, implicated in neurodegenerative disease and HIV/drug-induced neurotoxicity. **(D)** Flow cytometry analysis of HIV receptor expression on HEK293T, U937, and H80 cells. Cells were blocked with 2% BSA plus Fc block and co-stained for either CD4 and CXCR4 (APC anti-human CD4, BioLegend cat. no. 317416; PE anti-human CD184 \[CXCR4], BioLegend cat. no. 306505) or CD4 and CCR5 (PE anti-human CD4, BioLegend cat. no. 357403; APC/Cyanine7 anti-human CD195 \[CCR5], BioLegend cat. no. 359110). HEK293T cells (negative control) lacked CD4, CXCR4, and CCR5 expression, whereas U937 cells (positive control) expressed all three markers. H80 cells did not express CD4 or CCR5 but showed detectable CXCR4 expression. These findings suggest that H80 cells lack main HIV receptor CD4 and chemokine receptor CCR5 for HIV entry, despite expressing CXCR4.

Furthermore, to demonstrate that H80 cells possess the molecular characteristics of differentiated neurons, we focused on the expression of microtubule-associated protein 2 (MAP2). MAP2 is a neuron-specific cytoskeletal protein that is predominantly localized to dendrites, where it plays a pivotal role in stabilizing microtubules and maintaining neuronal morphology. MAP2 expression is therefore widely recognized as a hallmark of neuronal differentiation and identity. We initially assessed MAP2 expressions in H80 cells using immunofluorescence microscopy, with HEK293T cells serving as a non-neuronal reference (control), Microglial cells (unknown or known MAP2 negative cell line) and SH-SY5Y cells serving as neuronal reference (positive control or known to express MAP2). Notably, the expression of MAP2 was exclusively observed in H80 cells (**Figure 1B and supplementary S1**) and also in positive control (SH-SY5Y), where the protein displayed a distinct filamentous distribution throughout the cytoplasm, consistent with the structural organization seen in neurons. As anticipated, HEK293T cells lacked detectable MAP2 signal whereas SHS5Y has a strong MAP2 expression under identical staining conditions (**Figure 1B**). The expression of MAP2 indicates that H80 cells, but not HEK293T cells and microglial cells, belongs to neuronal lineage. To further corroborate these findings, we performed immunoblotting using total cellular lysates from H80 cells, HEK293T cells, and microglial cells. Consistent with the immunofluorescence results, MAP2 protein was robustly detected in H80 cell lysates, whereas it was undetectable in both HEK293T and microglial samples (**Figure 1C**). Importantly, the absence of MAP2 expression in microglial cells, another brain-resident glial cells of non-neuronal lineage, underscores the neuronal specificity of this marker. Together, these complementary assays provide convergent evidence that H80 cells exclusively express MAP2. The presence of MAP2 exclusively in H80 cells, but not in two distinct non-neuronal cell types, strongly supports the conclusion that H80 cells is a neuronal cell line that exhibits cytoskeletal and molecular features consistent with neuronal identity and differentiation.

Therefore, the co-expression of NeuN, MAP2, and Tau provides convergent and robust evidence that H80 cells exhibit hallmark neuronal features (**Figures 1A to C**, and **Supplementary Figure S1**). Collectively, these findings support the classification of H80 cells as a neuronal-like cell model. The presence of these canonical neuronal markers not only confirms their neuronal identity but also highlights their suitability as a versatile platform for investigating neuron-specific molecular mechanisms. In particular, H80 cells offer a valuable system for studying signaling pathways involved in the regulation of Tau protein phosphorylation and activity, as well as broader processes underlying neuronal function and neurotoxicity.

Since our study focuses on HIV induced neurotoxicity and neurons are not directly infected by HIV [17], we next examined the expression of the key HIV entry receptors and co-receptors in H80 cells (**Figure 1D**). To determine the expression profile of HIV entry receptors on H80 cells, we performed flow cytometric analysis for CD4, CCR5, and CXCR4. Cells were co-stained with either CD4 (APC anti-human CD4 from BioLegend cat no 317416) and CXCR4 (PE anti-human CD184 (CXCR4) Antibody from BioLegend cat no. 306505) or CD4 (PE anti-human CD4 Antibody from BioLegend cat no. 357403 and CCR5 (APC/Cyanine7 anti-human CD195 (CCR5) Antibody from BioLegend cat no. 359110). Our results demonstrated that H80 cells lack detectable CD4 expression under basal conditions, indicating the absence of the primary receptor required for productive HIV entry. Notably, approximately 20% of H80 cells expressed surface CXCR4, whereas CCR5 expression was undetectable. These findings suggest that while H80 cells are unlikely to support productive HIV infection due to the absence of CD4, the presence of CXCR4 on a subset of cells may render them responsive to HIV-associated proteins or signaling pathways, thereby contributing to HIV-induced neurotoxicity. To ensure assay specificity and reliability, HEK293T cells were included as a negative control and showed no detectable expression of CD4, CCR5, or CXCR4. In contrast, U937 cells served as a positive control and exhibited robust basal expression of CD4 and both co-receptors. Collectively, these data further support the suitability of H80 cells as a model to study HIV-mediated neurotoxic effects independent of direct viral infection.

Altogether, our findings demonstrate that H80 cells exhibit key neuronal characteristics, as evidenced by the expression of canonical neuronal markers, including NeuN, MAP2, and Tau. In addition, receptor profiling revealed that H80 cells lack detectable expression of CD4 and CCR5 but express the chemokine receptor CXCR4 on a subset of cells. Previous studies, including those by Kaul et al., have shown that both CXCR4 and CCR5 can mediate HIV-associated neuronal injury, while CCR5 may also engage neuroprotective signaling pathways [57]. The selective expression of CXCR4 in H80 cells is particularly noteworthy, as it suggests that analogous to neurons, these cells may be responsive to HIV-associated proteins and signaling events linked to CXCR4 engagement, despite the absence of productive viral entry. This receptor profile closely aligns with current understanding that neuronal damage in NeuroHIV is largely mediated indirectly through viral proteins and host signaling pathways rather than direct infection.

Based on these observations, we next sought to determine how HIV exposure influences neuronal stress responses and neurotoxicity in this model, with a particular focus on Tau phosphorylation, a well-established marker of tauopathy. Given the confirmed neuronal phenotype of H80 cells and their expression of CXCR4, these cells provide a biologically relevant system to study HIV-induced neuronal dysfunction independent of productive infection. Accordingly, we directly exposed H80 cells to HIV to investigate the underlying molecular mechanisms driving HIV-associated neurotoxicity and Tau dysregulation, enabling us to dissect signaling pathways that contribute to neurodegenerative processes in the context of NeuroHIV.

### HIV exposure upregulates RSK1 expression

To investigate the signaling pathways underlying HIV-induced tauopathy, H80 cells were exposed to HIV virions (strain 93/TH/051, R5- and X4-tropic virus, dual-tropic HIV-1). Because neurons lack the primary HIV receptor (CD4), they are resistant to productive infection; thus, analogous to neurons, this model allows us to specifically examine HIV-mediated signaling and neurotoxic effects independent of viral replication. HIV virions were generated by infecting Jurkat T cells, and successful infection was confirmed by immunoblot detection of the viral capsid protein p24 in infected cell lysates (**Figure 2A**). Virus-containing supernatants from either Jurkat or MT-4 cells were then collected and used to expose H80 cells using a two-round spinfection (spinoculation) protocol, which enhances viral contact and ensures efficient exposure of neuronal cells to viral particles. As a negative control, H80 cells were exposed to supernatants from uninfected Jurkat/MT-4 cells (**Figure 2A**). Following exposure, H80 cells were harvested at 24 and 48 hours after the second spinfection for downstream analyses. Total RNA and protein lysates were collected to assess changes in intracellular signaling pathways and Tau phosphorylation status, enabling us to define the molecular mechanisms by which HIV exposure induces neuronal stress and Tau dysregulation in this system.

**Figure 2:**
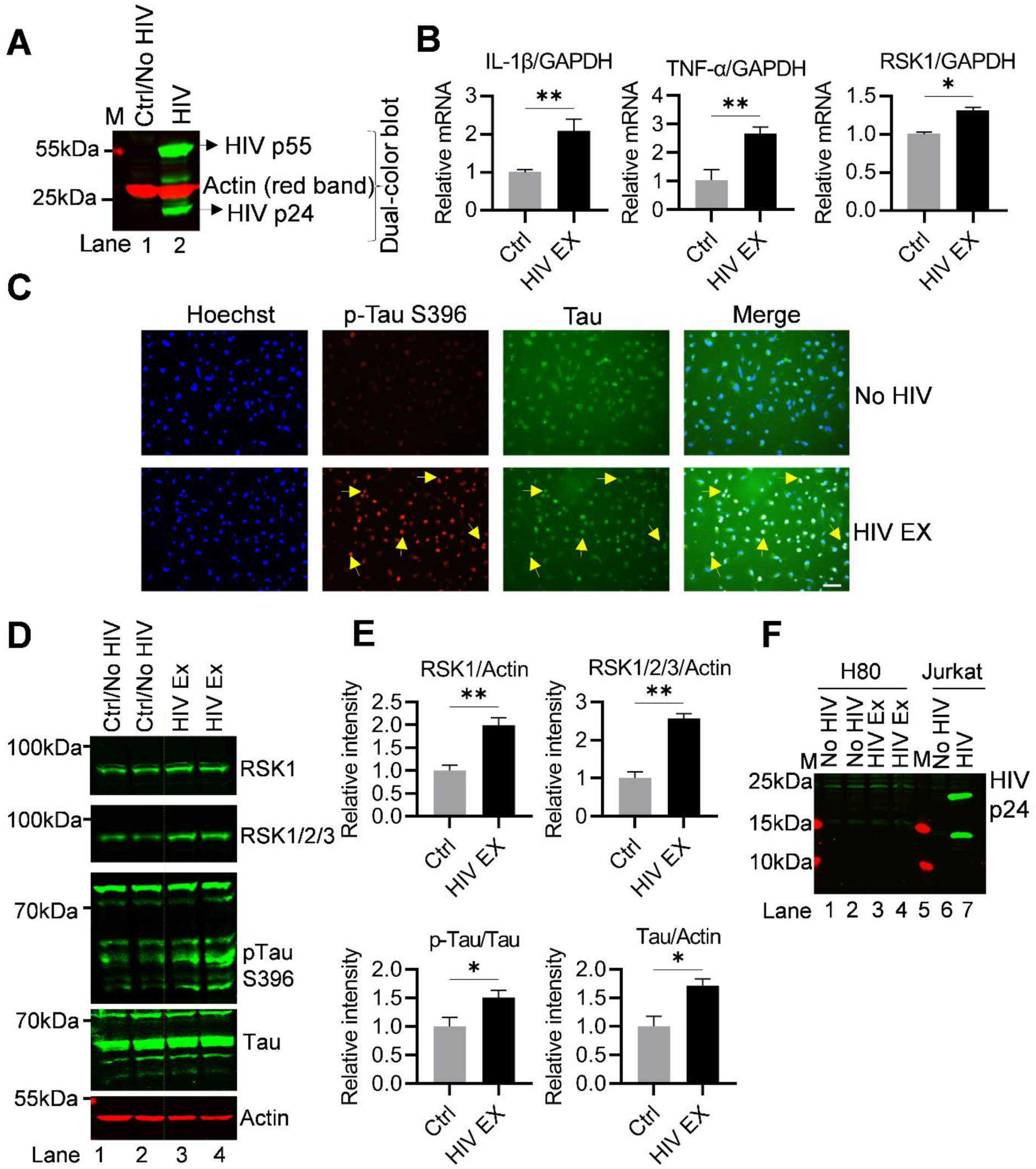
HIV exposure induces RSK1 upregulation and promotes Tau phosphorylation in H80 cells. **(A)** Immunoblot analysis of total cell lysates from Jurkat T cells infected with replication-competent HIV or uninfected (Ctrl), confirming HIV infection by detection of the HIV p24 capsid protein. Culture supernatants from uninfected Jurkat cells (control) and from HIV-infected cells (containing replication competent virion particles) were collected and used to expose H80 cells via spinoculation (2 h at 1,000 rpm). Cells were subsequently reseeded, and on the following day subjected to a second round of spinoculation with HIV virions under the same conditions, followed by seeding onto 100-mm plates. **(B)** Twenty-four hours post-exposure, total RNA was extracted from H80 cells and analyzed by reverse transcription–quantitative PCR (RT–qPCR) for the expression of IL-1β, TNF-α, and RSK1, relative normalized to GAPDH. HIV-exposed cells showed transcriptional upregulation of IL-1β, TNF-α, and RSK1 relative to No HIV controls. **(C)** After 48 h post-exposure, cells were subjected to immunofluorescence staining using antibodies against phosphorylated Tau at Ser396 and Tau. HIV exposure enhanced increased Tau phosphorylation compared to No HIV exposed controls. Yellow arrows indicate representative phosphorylation-positive sites in the immunofluorescence images. The scale bar represents 10 µm. **(D–E)** To validate these findings, H80 cells were cultured in four independent dishes (two biological replicates per condition), and whole cell lysates were collected 48 h after HIV exposure. Protein lysates were prepared separately from each dish and quantified. Equal amounts of protein were loaded and analyzed by immunoblotting for RSK1, RSK1/2/3, phosphorylated Tau (p-Tau-S396), and Tau, with actin or total protein as loading controls. Immunoblotting confirmed upregulation of RSK1, increased phosphorylation of Tau at Ser396, and a moderate elevation in Tau protein levels in HIV-exposed H80 cells. Densitometric analysis of immunoblots was performed by normalizing band intensities to β actin or total protein, with values expressed relative to the control (Ctrl/No HIV). **(F)** To assess whether H80 cells were susceptible to HIV infection, lysates from HIV–exposed and No HIV exposed H80 cells were examined for HIV p24 by immunoblotting, alongside Jurkat T cells included as positive (infected) and negative (uninfected) controls. Immunoblots are representative of at least three independent biological replicates. Data are presented as mean ± S.D. Statistical significance was assessed using an unpaired, two tailed Student’s t test. Significance is indicated as P < 0.05 (*) and P < 0.01 (**).

To assess the impact of HIV exposure on the cellular transcriptional machinery and to confirm successful viral exposure on H80 cells, we quantified mRNA expression levels of representative inflammatory and signaling genes using qRT-PCR at 24 hours post-exposure. HIV-exposed H80 cells expressed a robust upregulation of interleukin-1β (IL-1β) and tumor necrosis factor-alpha (TNF-α) transcripts, both of which are central mediators of proinflammatory signaling cascades (**Figure 2B**). Consistent with previous reports demonstrating that neurons can produce cytokines in response to HIV-associated stress [58], this pronounced induction provides strong evidence of effective HIV exposure and activation of innate immune signaling in H80 cells. In addition, we observed a modest but reproducible increase in RSK1 mRNA expression. Given the established role of RSK1 as a downstream effector of MAPK signaling, this finding suggests early activation of stress-responsive and pathogen-associated signaling pathways, complementing the observed inflammatory response. Collectively, these transcriptional changes not only confirmed the upregulation of RSK1 upon HIV exposure but also underscore the functional reactivation of the cells, thereby validating the functional responsiveness of H80 cells to HIV exposure. These results further strengthen the biological relevance of our H80 neuronal model for dissecting the molecular mechanisms underlying HIV-induced neuroinflammation, stress signaling, and Tau-associated pathology.

### HIV exposure-driven RSK1 upregulation induces Tau phosphorylation

To investigate whether HIV exposure promotes pathological Tau modification, we first performed immunofluorescence staining using an antibody specific for the phosphorylated Tau protein at Ser396 (p-Tau-Ser396), a site commonly associated with neurotoxicity and neurodegeneration. Compared with controls (exposing without the virus), HIV-exposed H80 cells displayed a marked increase in p-Tau-Ser396 signal, suggesting that HIV exposure drives Tau phosphorylation (**Figure 2C**). Notably, in addition to the phosphorylation of Tau, HIV exposure markedly upregulates RSK1 (**Supplementary Figure S2**). The concurrent induction of RSK1 suggested a role of this kinase in promoting Tau phosphorylation. These findings indicate that the effect of HIV exposure on Tau phosphorylation facilitated through the regulation of RSK1, thereby highlighting a mechanistic link between RSK1 expression and Tau phosphorylation.

To substantiate these observations, we conducted immunoblot analysis of whole-cell lysates following HIV exposure (under the same conditions). H80 cells were cultured in four independent dishes (two biological replicates per condition), and whole cell lysates were collected 48 h after HIV exposure. Protein lysates from each dish were prepared and quantified individually. Equivalent amounts of protein were resolved by immunoblotting to assess RSK1, RSK1/2/3, phosphorylated Tau (p Tau S396), and Tau, using actin or total protein as loading controls. Consistent with the immunofluorescence imaging results, immunoblotting revealed a robust induction of p-Tau-Ser396 in HIV-exposed cells (lanes 3-4) compared to control (exposing with supernatant from uninfected cells, lanes 1-2). In contrast, total Tau protein levels exhibited only a modest increase, indicating that HIV exposure predominantly induces post-translational modification of Tau, rather than significantly altering its overall expression. Notably, this increase in Tau phosphorylation was accompanied by a pronounced upregulation of RSK1, as well as elevated levels of the RSK1/2/3 isoforms, suggesting activation of the RSK signaling pathway in response to HIV exposure. These findings indicate that RSK1 activation occurs in parallel with HIV-induced Tau phosphorylation and may serve as a critical molecular link between viral exposure and Tau dysregulation in H80 cells (**Figure 2D**). Quantitative densitometric analysis of the immunoblot signals, normalized to β actin and/or total protein, revealed a statistically significant increase in the levels of both RSK1 and p-Tau Ser396 in HIV exposed H80 cells relative to controls/No HIV (**Figure 2E**). To further assess whether H80 cells support productive HIV infection, lysates from HIV-exposed and control (no HIV) conditions were analyzed by immunoblotting using an anti-HIV p24 antibody, with Jurkat T cells included as positive (HIV-exposed) and negative (uninfected) controls (**Figure 2F**). Consistent with the absence of CD4 expression, p24 was not detected in H80 cell lysates, indicating that these cells do not support productive HIV infection and are instead exposed to HIV virions without undergoing viral replication. Importantly, despite the lack of productive infection, exposure to HIV virions was sufficient to induce robust activation of RSK1 signaling in H80 cells, which correlated with increased Tau phosphorylation. Given the well-established roles of RSK1 in nuclear gene regulation and cytoplasmic signaling crosstalk, these findings suggest that HIV-induced activation of RSK1 serves as a key mediator linking viral exposure to downstream neuronal stress responses. Collectively, our results support a model in which HIV virion exposure, independent of productive infection, triggers RSK1 activation, leading to pathological Tau phosphorylation and tauopathy-associated signaling, thereby establishing a mechanistic connection between HIV exposure and neuronal dysfunction mediated through the RSK1 pathway.

### Cocaine enhances Tau phosphorylation by upregulating and activating RSK1

To investigate whether cocaine contributes to Tau pathology, we chronically exposed H80 cells to cocaine twice daily for 2 days and subsequently assessed Tau phosphorylation by immunofluorescence (**Figure 3A**). Cocaine-exposed H80 cells exhibited a pronounced increase in phosphorylated Tau (p-Tau-Ser396) compared with untreated controls, while the total abundance of Tau protein remained unchanged (**Figure 3B**). These findings indicate that cocaine primarily promotes post-translational modification of Tau, rather than significantly altering its overall expression. Given that our previous findings demonstrated that HIV exposure upregulates RSK1 (**Figure 2**), we next examined whether cocaine-induced Tau phosphorylation is similarly associated with upregulation in RSK1 expression. Immunofluorescence analysis revealed a significant upregulation of RSK1 following chronic cocaine exposure (**Supplementary Figure S2**). Thus, parallel increment in RSK1 expression and Tau phosphorylation suggested crucial role for RSK1 signaling in mediating cocaine-induced Tau modification. These findings were further validated in an independent experiment, confirming the reproducibility of RSK1 upregulation and enhanced Tau phosphorylation in response to cocaine exposure in H80 cells (**Figure 3C–3F**).

**Figure 3:**
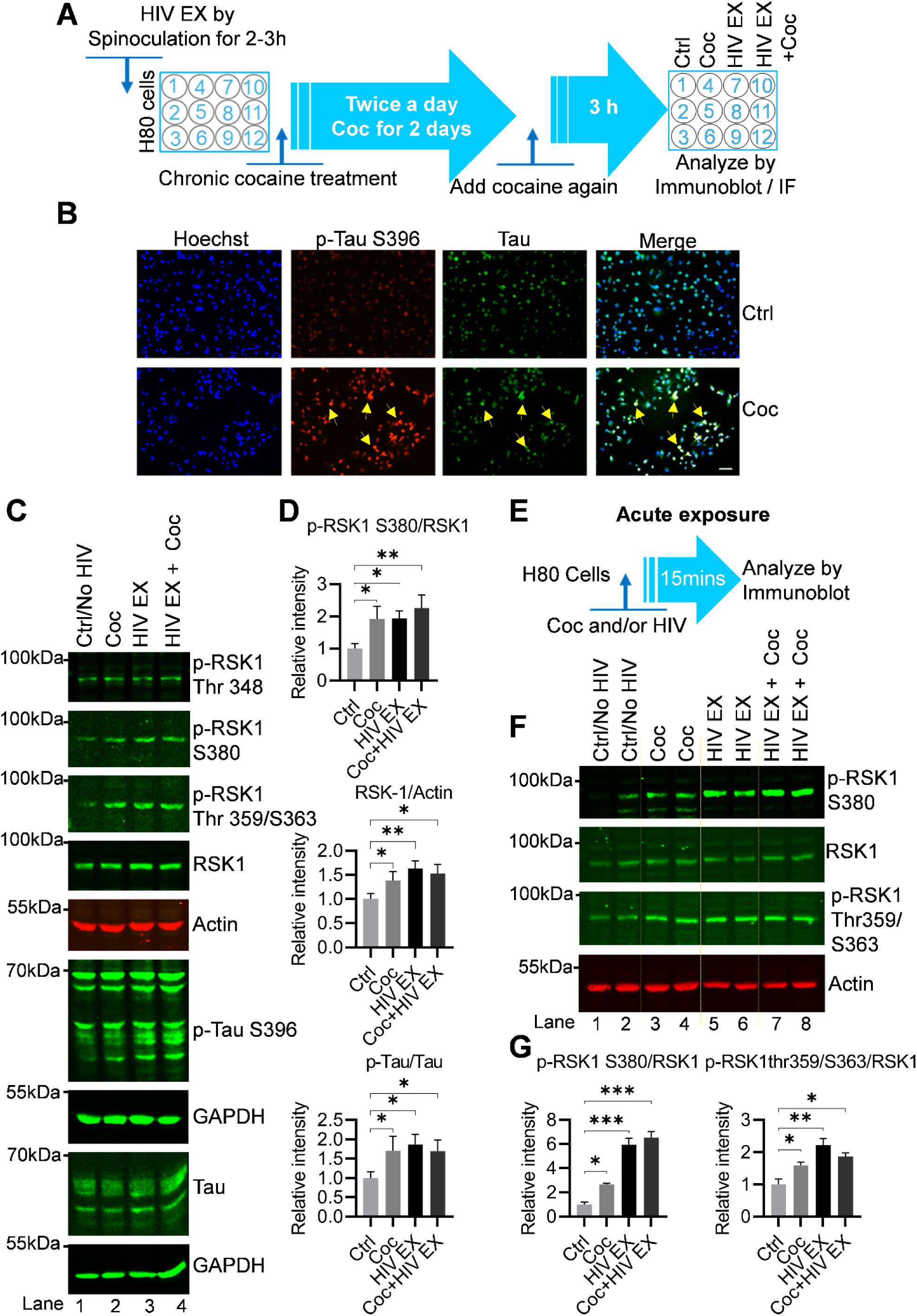
Cocaine activates and upregulates RSK1 and promotes Tau phosphorylation in H80 cells. (A) Schematic representation of the protocol for the IF and Immunoblot assay detailing treatment with the chronic cocaine and HIV exposure **(B**) Immunofluorescence analysis of H80 cells chronically exposed to cocaine (twice daily for 2 days) revealed a marked increase in phosphorylated Tau at Ser396 (p-Tau-Ser396) compared with untreated controls, while Tau levels remained unchanged, indicating that cocaine enhances Tau phosphorylation without altering overall Tau protein. Yellow arrows indicate representative phosphorylation-positive sites in the immunofluorescence images. The scale bar represents 10 µm. (**C–D**) Immunoblot analysis of whole-cell lysates from H80 cells treated with cocaine, HIV, or both revealed that cocaine modestly increased total RSK1 expression and its phosphorylation at Ser380, a marker of catalytic activation, while minimally affecting Thr348 phosphorylation. HIV exposure produced a robust increase in total RSK1 and phosphorylation at multiple sites (Ser380, Thr348, Thr359, and Ser363), far exceeding the effects of cocaine alone. Both cocaine and HIV significantly elevated p-Tau-Ser396 relative to untreated controls, with HIV exerting a stronger effect. Co-exposure to cocaine and HIV resulted in a higher, though not strictly additive, increase in Tau phosphorylation, suggesting convergence on overlapping signaling pathways. Densitometric analysis of immunoblots was performed by normalizing band intensities to β actin or total protein, with values expressed relative to the control (ctrl/No HIV). **(E)** Schematic representation of the protocol for immunoblots with acute cocaine and HIV exposure. **(F-G)** H80 cells of different passages were cultured in eight independent dishes (two biological replicates per condition), and whole–cell lysates were collected 15 min after exposure to cocaine, HIV, or the combined treatment. Protein lysates were prepared separately from each dish and quantified. Equal amounts of protein were loaded and analyzed by immunoblotting for p-RSK1 S380, p-RSK1 Thr359 and S363, and RSK1, with actin as loading controls. Immunoblotting confirmed activation of RSK1, increased phosphorylation of RSK1 at S380, Thr359 and S363, in HIV-exposed H80 cells. Densitometric analysis of immunoblots was performed by normalizing band intensities to RSK1, with values expressed relative to the control (Ctrl/No HIV). Immunoblots are representative of at least three independent biological replicates. Data are presented as mean ± S.D. Statistical significance was assessed using one way ANOVA with Dunnett’s multiple comparisons test. Significance is indicated as P < 0.05 (*) and P < 0.01 (**). Together, these findings implicate RSK1 activation as a key mediator of cocaine- and HIV-driven Tau phosphorylation, highlighting a shared molecular axis underlying neurodegenerative processes in the context of substance use and HIV exposure.

To further solidify the data obtained through immunofluorescence analysis, we chronically treated H80 cells alone or in combination with cocaine and HIV as shown in **Figure 3A**. The cell lysate was analyzed by Immunoblotting. The immunoblotting of whole-cell lysates revealed that cocaine produced a modest but reproducible increase in total RSK1 protein, accompanied by an elevation in its phosphorylation at Thr 348, Thr 359, S363 and S380, which marks functionally active form of RSK1. Given that specific posttranslational modification of RSK1 was quantitatively correlated with the increase in p-Tau-Ser396, implicating RSK1 as a mediator of cocaine-driven Tau modification (**Figure 3C and 3D**). Both cocaine and HIV significantly increased p-Tau-Ser396 relative to controls, although the magnitude of the effect differed: HIV produced a robust elevation in Tau phosphorylation, whereas cocaine induced a more modest increase. Notably, co-exposure to cocaine and HIV did not result in strictly additive effects (but it is more on higher side), suggesting that these stimuli converge on overlapping molecular pathways. Analysis of RSK1 activation under these conditions revealed that HIV strongly enhanced both total RSK1 expression and its phosphorylation, including at S380, Thr348, Thr359/ and S363 far exceeding the effects of cocaine alone. HIV exposure also led to a moderate increase in Thr348 phosphorylation, further distinguishing its mode of RSK1 regulation from cocaine. Importantly, the degree of RSK1 activation in each condition closely aligned with the extent of Tau phosphorylation, strengthening the link between RSK1 signaling and Tau modification. Together, these results establish that both cocaine and HIV drive activation of the RSK1 signaling axis in H80 cells, albeit with distinct magnitudes and mechanistic profiles (**Figure 3C and 3D**). Cocaine modestly increases RSK1 expression and Tau phosphorylation, whereas HIV produces a more potent and broader activation of RSK1, resulting in stronger downstream Tau modification. These findings provide mechanistic insight into how viral infection and substance use converge on a shared signaling pathway to promote Tau pathology.

To delineate the immediate signaling responses triggered by cocaine and HIV, we examined the acute activation of RSK1 in H80 neuronal cells following short term exposure to each stimulus. H80 cells were cultured in eight independent dishes, providing two biological replicates for each of the four treatment conditions (Control, cocaine, HIV, and cocaine + HIV). Cells were exposed acutely for 15 minutes (**Figure 3E**), after which they were harvested and lysed. Protein lysates from each dish were prepared and quantified individually, and equivalent amounts of total protein were resolved by immunoblotting to evaluate phosphorylation of RSK1 at the activating sites Ser380 and Thr359/Ser363, alongside total RSK1 levels, using actin or total protein as loading controls. Acute exposure to either cocaine or HIV elicited rapid and robust activation of the RSK1 signaling pathway in H80 cells. Immunoblot analysis revealed marked increases in RSK1 phosphorylation at both Ser380 and Thr359/Ser363 relative to controls (**Figure 3F and 3G**). This activation was consistently observed across biological replicate lanes corresponding to each treatment group (lanes 3–4, 5–6, and 7–8 compared with lanes 1–2), demonstrating strong reproducibility of these rapid phosphorylation events. Notably, the magnitude of RSK1 activation differed between stimuli. While cocaine induced a clear and reproducible increase in phosphorylation at both regulatory sites (p-RSK-1 Thr359/S363 and S380), HIV exposure elicited a substantially stronger response, producing the highest levels of RSK1 activation among all acute treatment conditions.

Together, these findings establish that RSK1 is a rapidly responsive kinase activated within minutes of cocaine or HIV exposure, and that the amplitude of this response is greater under HIV stimulation than under cocaine alone. This further reinforces the pattern observed under chronic exposure conditions, underscoring the consistency of cocaine and HIV–driven enhancement of RSK–1 activation across temporal paradigms, reinforcing its central role as an early signaling mediator linking cocaine and HIV exposure to downstream neuronal stress responses and Tau pathology.

### HIV or Cocaine converge on GSK3β inhibition through Ser9 phosphorylation

GSK3β is a key regulator of neuronal signaling and a well-established mediator of Tau phosphorylation [59–63]. The enzymatic activity of GSK3β is primarily controlled through its phosphorylation at the serine-9 (Ser9) residue, (p-GSK3β-Ser9), which serves as an inhibitory modification that limits substrate access and thereby inhibits GSK3β kinase activity. Given the well-established role of GSK3β in mediating Tau pathology, we next examined whether HIV and cocaine influence the regulation of this kinase. As shown in **Figures 2 and 3**, both HIV and cocaine exposure markedly enhanced Tau phosphorylation, raising the possibility that these effects are mediated, at least in part, through altered GSK3β activity. To investigate further, we specifically assessed the phosphorylation status of GSK3β at its inhibitory Ser9 residue, thereby determining whether HIV and cocaine relieve the inhibitory regulation of GSK3β and contribute to the observed increase in Tau phosphorylation. Using total cellular lysates from **Figure 2A**, we examined the phosphorylation status of GSK3β at the inhibitory Ser9 site by immunoblotting. Surprisingly, our results demonstrated a marked increase in Ser9 phosphorylation in response to HIV infection in Jurkat cells, while the levels of total GSK3β remained unchanged (**Figure 4A**). This increase in inhibitory phosphorylation suggests that GSK3β becomes inactivated upon HIV infection or under conditions of ongoing viral infection.

**Figure 4:**
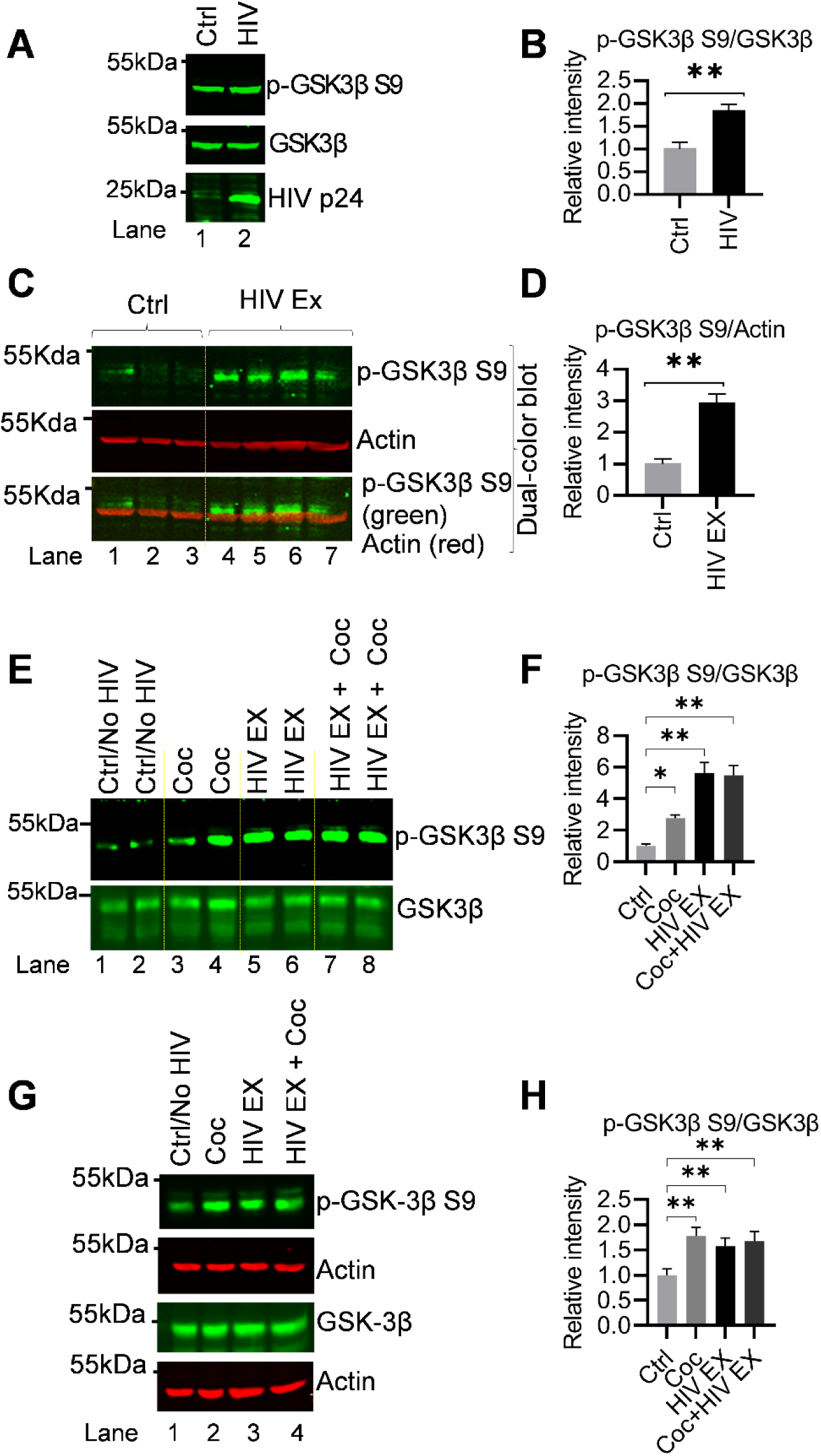
HIV and cocaine converge on GSK3β inactivation via Ser9 phosphorylation. (A-B) Immunoblot analysis of Jurkat T cells infected with HIV or uninfected (control) revealed a marked increase in GSK3β phosphorylation at the inhibitory Ser9 site (p-GSK3β-Ser9) upon HIV infection, while GSK3β levels remained unchanged, indicating functional inactivation of GSK3β during viral infection in immune cells. **(C-D)** Acute exposure of H80 cells (15 min) to supernatants from HIV-infected Jurkat cells induced a pronounced increase in Ser9 phosphorylation compared to control (uninfected) supernatant, demonstrating that HIV exposure rapidly modulates host kinase signaling even in non-permissive cells lacking CD4. **(E-F)** H80 cells were cultured in eight independent dishes (two biological replicates) under each identical condition and whole-cell lysates were collected 15 min after cocaine, HIV exposure and cocaine plus HIV exposure. Protein lysates were prepared separately from each dish and quantified. Equal amounts of protein were loaded and analyzed by immunoblotting for p-GSK3β S9, and GSK3β. Immunoblotting confirmed inactivation of GSK3β, increased phosphorylation of GSK3β at S9, in cocaine, HIV exposed and cocaine along with HIV-exposed H80 cells compared to Ctrl/No HIV. Densitometric analysis of immunoblots was performed by normalizing band intensities to GSK3β, with values expressed relative to the control (Ctrl/No HIV). Immunoblots are representative of at least three independent biological replicates. **(G-H)** Immunoblot analysis of H80 cells exposed for 48 h to cocaine (chronic exposure), HIV virions, or both (Schematic representation in Figure 3A) revealed robust Ser9 phosphorylation under all conditions, while GSK3β levels remained constant, confirming post-translational regulation rather than changes in protein abundance. Combined exposure to HIV and cocaine produced an inhibitory effect of GSK3β similar to individual treatments. Densitometric analysis of immunoblots was performed by normalizing band intensities to GSK3β, with values expressed relative to the control (Ctrl/No HIV). Immunoblots are representative of at least three independent biological replicates. Data are presented as mean ± S.D. Statistical significance was assessed using an unpaired, two tailed Student’s t test (for B and D) or one way ANOVA with Dunnett’s multiple comparisons test (for F and H). Significance is indicated as P < 0.05 (*) and P < 0.01 (**).

To determine whether HIV exposure directly modulates GSK3β activity, we exposed H80 cells for 15 min to the supernatant derived from HIV-infected cells (Jurkat infected with HIV), while supernatant from uninfected cell cultures (Jurkat uninfected with HIV) was used as a control, as shown in **Figure 2A**. After 15 min exposure, the cells were harvested, and protein lysates were subjected to immunoblot analysis to evaluate the phosphorylation status of GSK3β at Ser9 site, marking functionally inactive form of GSK3β. We observed a pronounced increase in Ser9 phosphorylation of GSK3β in HIV-exposed cells compared to the control (**Figure 4B and 4C**). Since phosphorylation at Ser9 is known to suppress GSK3β catalytic activity, this increase strongly suggests that acute HIV exposure enhance posttranslational modification of GSK3β at Se9, which functionally inactivates GSK3β. These findings indicate that HIV exposure can rapidly influence host kinase signaling pathways.

Subsequently, we examined how acute exposure to cocaine and HIV modulates GSK3β signaling in H80 cells. To evaluate these rapid effects, cells were cultured in eight independent dishes, providing two biological replicates per treatment condition, and exposed for 15 minutes to cocaine, HIV, or both. Following treatment, cells were harvested and lysed, and protein lysates from each dish were prepared and quantified individually. Equal amounts of total protein were subjected to immunoblot analysis to assess phosphorylation of GSK3β at Ser9, with total protein levels serving as loading controls. Both cocaine and HIV independently elicited a clear increase in Ser9 phosphorylation of GSK3β (Lane 3-4, lane 5-6 and lane 7-8 compared to lane 1-2), indicating enhanced inhibitory modification of the kinase (**Figure 4E and 4F**). Notably, the consistent elevation of Ser9 phosphorylation in HIV exposed samples further confirms the reproducibility of HIV-mediated GSK3β inactivation, as also observed in **Figures 4B and 4C**. These findings demonstrate that acute exposure to either cocaine or HIV is sufficient to rapidly inactivate GSK3β, revealing a shared regulatory mechanism by which both stimuli attenuate GSK3β activity in H80 cells. Importantly, this occurs despite the observed increase in Tau phosphorylation, further supporting the notion that cocaine- and HIV-induced Tau dysregulation proceeds through GSK3β-independent signaling pathways, likely involving alternative kinases such as RSK1.

To further substantiate our finding that both HIV and cocaine are independently able to inactivate GSK3β, we performed immunoblot analysis to assess inhibitory phosphorylation of GSK3β at Ser9. H80 cells were chronically exposed for 48 hours to cocaine, HIV virions, or a combination of both. Under each condition, cocaine alone, HIV alone, or combined HIV plus cocaine exposure, we observed a robust increase in Ser9 phosphorylation relative to untreated controls, whereas total GSK3β protein levels remained unchanged. These results indicate that the effects of HIV and cocaine on GSK3β are mediated through post-translational inhibitory modification, rather than changes in protein abundance. Notably, combined exposure to HIV and cocaine also produced a clear increase in p-GSK3β Ser9 compared with control cells (**Figure 4G and H**), confirming that both stimuli converge on functional inactivation of GSK3β. The consistency of this response across all treatment groups strongly supports the conclusion that HIV and cocaine each suppress GSK3β activity in H80 neuronal cells. This finding is particularly striking because GSK3β is widely recognized as a major Tau kinase. Accordingly, if GSK3β is the main Tau kinase, inactivation of GSK3β would be expected to reduce Tau phosphorylation. In contrast, we observed the opposite outcome: despite clear evidence of GSK3β inactivation, Tau phosphorylation was markedly increased following exposure to HIV and/or cocaine (**Figures 2 and 3**). This apparent paradox strongly suggests that Tau phosphorylation in this setting is driven through an alternative pathway that is independent of GSK3β activity. Our data point towards RSK1 as a likely upstream mediator of this effect. Indeed, both HIV and cocaine induced significant upregulation and activation of RSK1, coinciding with enhanced Tau phosphorylation under conditions in which GSK3β remained inactivated. These observations support a model in which RSK1-driven signaling bypasses the need for active GSK3β and sustains pathological Tau phosphorylation, thereby promoting tauopathy-associated neuronal stress responses.

Collectively, these findings demonstrate that HIV and cocaine independently converge on GSK3β inactivation via Ser9 phosphorylation yet simultaneously promote Tau hyperphosphorylation through a distinct upstream mechanism, most likely involving RSK1. This convergence on inhibitory GSK3β signaling, together with activation of an alternative Tau-phosphorylating pathway, identifies a critical molecular axis by which viral exposure and substance use disrupt neuronal signaling, ultimately contributing to Tau dysregulation and the CNS impairments, including neuropathological processes associated with HAND.

### Cocaine, but not HIV exposure, activates AKT1 signaling through phosphorylation at Thr308 and Ser473

Phosphorylation of GSK3β at Ser9, a critical inhibitory modification, is tightly regulated by upstream kinases, most notably AKT, which plays a central role in neuronal signaling cascades and survival pathways [34]. AKT-mediated phosphorylation of GSK3β at Ser9 serves as a key inhibitory checkpoint that suppresses GSK3β catalytic activity and prevents excessive substrate phosphorylation. Given our findings that both HIV and cocaine independently increase Ser9 phosphorylation on GSK3β, thereby promoting its inactivation, we next sought to determine whether these are direct effect are mediated through activation of the AKT signaling pathway. To investigate this, we examined the phosphorylation status of AKT at its regulatory sites that control its functional activity. We hypothesized that stimulation of AKT activity would restrict GSK3β activity by catalyzing its phosphorylation at Ser9 (p-GSK3β-Ser9). This approach allowed us to directly evaluate whether HIV and cocaine converge upstream on AKT to regulate GSK3β activity and thus, contribute to Tau hyperphosphorylation.

To determine the regulation of AKT signaling pathway upon HIV and cocaine exposure, we performed immunofluorescence staining for phosphorylated AKT at Ser473 (p-AKT-Ser473), a well-established marker of AKT activation and a critical modification required for full kinase activity. H80 cells were chronically exposed for 2 days to cocaine, HIV virions, or both, after which AKT1 phosphorylation status was evaluated (**Figure 5A**). Cocaine-treated cells displayed a robust, reproducible and significant increase in p-AKT-Ser473 fluorescence compared with untreated controls, indicating that cocaine strongly activates AKT signaling pathway (**Figure 5B**). In contrast, HIV exposure alone did not show any effect in p-AKT-Ser473 levels under the same conditions (**Supplementary Figure S3**), suggesting that HIV does not directly induce AKT activation in this context. Notably, total AKT protein levels were unaffected across all conditions, confirming that the observed changes were attributable to post-translational regulation of phosphorylation rather than changes in protein expression or stability.

**Figure 5:**
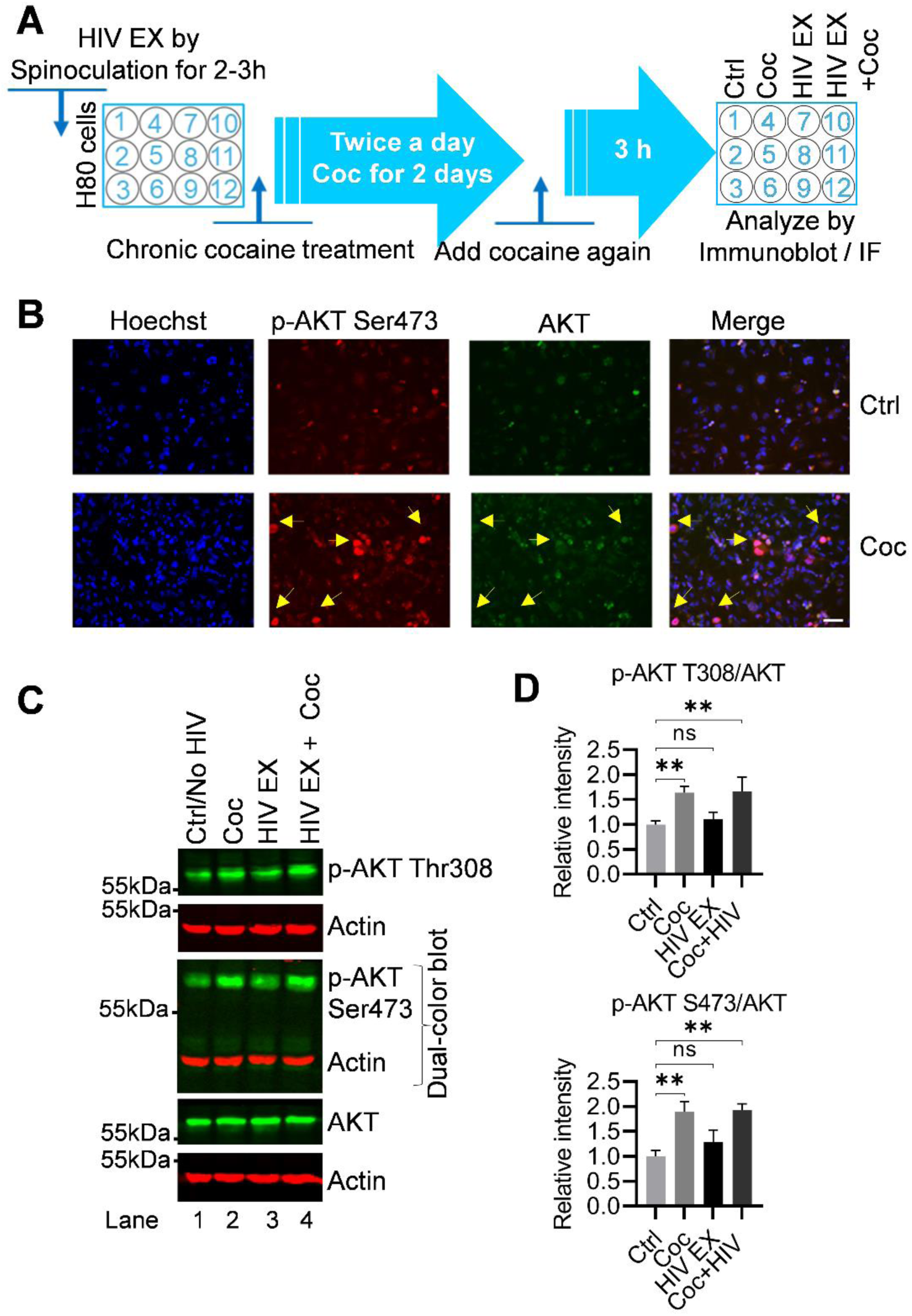
Cocaine activates AKT signaling in H80 cells, whereas HIV exposure does not activate AKT. (**A**) Schematic representation of the protocol for the IF and Immunoblot assay detailing treatment with the chronic cocaine and HIV exposure **(B)** Immunofluorescence analysis of H80 cells chronically exposed to cocaine (twice daily for 2 days) revealed a robust increase in phosphorylated AKT at Ser473 (p-AKT-Ser473) compared with untreated controls, indicating strong activation of the AKT signaling pathway. Hoechst staining was used for nuclear visualization. AKT levels remained unchanged, confirming that the observed effect reflects post-translational regulation rather than changes in protein abundance. Yellow arrows indicate representative phosphorylation-positive sites in the immunofluorescence images. The scale bar represents 10 µm. HIV exposure alone did not alter p-AKT-Ser473 levels under the same conditions (data in supplementary). **(C)** Immunoblot analysis of whole-cell lysates from H80 cells exposed to cocaine, HIV, or both for 48 h demonstrated that cocaine significantly increased phosphorylation of AKT at both Thr308 and Ser473, modifications essential for full kinase activation. HIV exposure alone did not affect AKT phosphorylation, while combined treatment mirrored the effect of cocaine alone, indicating that cocaine exerts a dominant influence on AKT activation. **(D)** Densitometric quantification confirmed a significant increase in AKT phosphorylation at Thr308 and Ser473 in cocaine-treated and HIV+cocaine-treated cells, whereas HIV alone had no measurable impact. AKT protein levels remained constant across all conditions. Densitometric analysis of immunoblots was performed by normalizing band intensities to AKT, with values expressed relative to the control (Ctrl/No HIV). Immunoblots are representative of at least three independent biological replicates. Data are presented as mean ± S.D. Statistical significance was assessed using one way ANOVA with Dunnett’s multiple comparisons test. Significance is indicated as P < 0.01 (**) and ns denotes not significant.

To further validate our data obtained through immunofluorescence analyses and confirm the diverse impact of cocaine and HIV on AKT signaling, we performed immunoblot analysis using cell lysates from H80 cells following 2 days of chronic exposure to cocaine, HIV, or a combination of both and evaluate both the phosphorylation sites of AKT. Immunoblotting revealed that cocaine treatment induces a significant increase in phosphorylation of AKT at both Thr308 and Ser473 compared with untreated controls (**Figure 5C**). Since phosphorylation at Thr308 and Ser473 are both essential for full activation of AKT, the simultaneous increase in phosphorylation at these two regulatory sites strongly confirms that cocaine induces a robust activation of the AKT signaling pathway. In contrast, HIV exposure alone did not alter phosphorylation at either site (further validating our immunofluorescence results), demonstrating that HIV exposure alone does not enhance or activate AKT signaling pathway. Notably, combined treatment with cocaine and HIV reproduced the increase in phosphorylation pattern of AKT observed with cocaine alone, indicating that cocaine exerts a dominant stimulus in activating AKT signaling, even in the presence of viral exposure. This suggests that cocaine overrides any potential influence of HIV on this pathway. Quantitative densitometric analysis (**Figure 5D**) provided further support, showing a significant increase in AKT phosphorylation at both Thr308 and Ser473 in cocaine alone and HIV+ cocaine-exposed cells, while HIV exposure alone had no measurable impact relative to controls. Importantly, total AKT protein levels remained constant across all conditions, confirming that the observed changes reflect post-translational modifications rather than at gene expression. Together, these results establish that cocaine, but not HIV, selectively activates AKT signaling in H80 cells, with cocaine driving strong phosphorylation of AKT at both activation sites and dominating over HIV when both stimuli are present.

Collectively, these findings establish cocaine, but not HIV, as a potent activator of the AKT signaling pathway in H80 cells. Mechanistically, cocaine promotes sustained AKT phosphorylation at Thr308 and Ser473, which subsequently promotes inactivation of GSK3β by catalyzing its phosphorylation at Ser9 (p-GSK3β-Ser9). Thus, cocaine promotes GSK3β inactivation (p-GSK3β-Ser9) both directly through AKT stimulation and also via RSK1 activation. In contrast, HIV inactivates GSK3β exclusively through an AKT-independent mechanisms, primarily through RSK1 signaling.

Thus, cocaine and HIV converge on a shared downstream effector, GSK3β inactivation, but diverge in their upstream regulatory pathways. Cocaine acts through an AKT-dependent pathway that also involves RSK1 stimulation, whereas HIV acts predominantly through an AKT-independent, RSK1-driven pathway. This dual convergence and divergence highlight the complexity of signaling networks regulating Tau phosphorylation and underscore how viral exposure and substance use engage distinct yet overlapping molecular pathways to drive neurotoxicity and tauopathy.

### HIV and Cocaine upregulate RSK1 to drive Tau phosphorylation through a GSK3β-independent mechanism

Our findings thus far indicate that HIV and cocaine share a common upstream signaling pathway through the upregulation of RSK1 (**Figure 2 and 3**). However, their downstream signaling pathways diverge in a stimulus-specific manner. Notably, HIV does not activate the AKT signaling pathway (**Figure 5**), whereas cocaine exposure leads to robust AKT activation, as evidenced by increased phosphorylation at both key regulatory sites, Thr308 and Ser473 (**Figure 5A-D**). Despite this divergence, both stimuli ultimately converge on the inactivation of GSK3β, as demonstrated by increased inhibitory phosphorylation at Ser9 (**Figure 4**). This shared downstream effect establishes GSK3β as a critical point of signaling convergence. Importantly, this convergence results in a common pathological outcome, the enhanced Tau phosphorylation of Tau (**Figure 2 and 3**), a hallmark of neurodegenerative processes and tauopathy.

These observations suggest a model in which distinct upstream signaling pathways (AKT-dependent for cocaine and AKT-independent for HIV) converge on shared downstream nodes, while simultaneously engaging alternative kinase pathways, such as RSK1, to drive Tau phosphorylation. The persistence of Tau phosphorylation despite GSK3β inactivation further underscores the involvement of GSK3β-independent mechanisms, likely mediated by RSK1. Based on these findings, we next sought to delineate the relative contributions and mechanistic interplay between RSK1 and GSK3β in mediating Tau phosphorylation following HIV and cocaine exposure. This approach aims to clarify how these signaling pathways integrate to produce a shared neurotoxic phenotype, thereby providing deeper insight into the mechanistic basis of HIV-and substance use-mediated tauopathy and neurodegeneration.

To validate our findings and further define the effects of cocaine and HIV on Tau phosphorylation, we performed a comprehensive immunoblot analysis using lysates from H80 cells following 48 hours of chronic exposure to cocaine, HIV, or their combination. Cells were seeded into 12 independent culture dishes across ≥3 passages/days, yielding three biological replicates per condition (Control, cocaine, HIV, and cocaine + HIV). After treatment, cells from each dish were harvested and lysed individually, and equal amounts of total protein were subjected to immunoblot analysis. Consistent with our earlier observations (**Figures 2 and 3**), immunoblotting revealed that cocaine, HIV, and their combination each induced a significant increase in Tau phosphorylation at Ser396 compared with untreated controls (**Figures 6A and 6B**). These results robustly confirm and extend our previous findings, demonstrating that both stimuli, independently and in combination, promote sustained Tau hyperphosphorylation under chronic exposure conditions. Phosphorylation of Tau at Ser396 is widely recognized as a marker of pathological Tau, and the observed increase at this site strongly confirms that cocaine exposure, HIV exposure, and their combination each induce Tau hyperphosphorylation. In parallel, we assessed the activity of GSK3β under these conditions. Interestingly, despite the increase in Tau phosphorylation, GSK3β was found to be functionally inactivated, as evidenced by enhanced phosphorylation at its inhibitory Ser9 residue (**Figures 6A and 6B**). These findings demonstrate that both HIV virion exposure and chronic cocaine treatment promote Tau phosphorylation while simultaneously restricting GSK3β activity, indicating that Tau hyperphosphorylation occurs through a GSK3β-independent mechanism. Importantly, although both stimuli converge on GSK3β inactivation, they do so via distinct upstream pathways. HIV induces GSK3β inactivation in an AKT-independent manner, whereas cocaine mediates this effect through AKT activation, as reflected by increased phosphorylation at both Thr308 and Ser473. Together, these results highlight a critical mechanistic distinction in upstream signaling while reinforcing a shared downstream outcome, Tau hyperphosphorylation despite GSK3β inhibition, suggesting the involvement of alternative kinases, such as RSK1, in driving Tau pathology.

**Figure 6:**
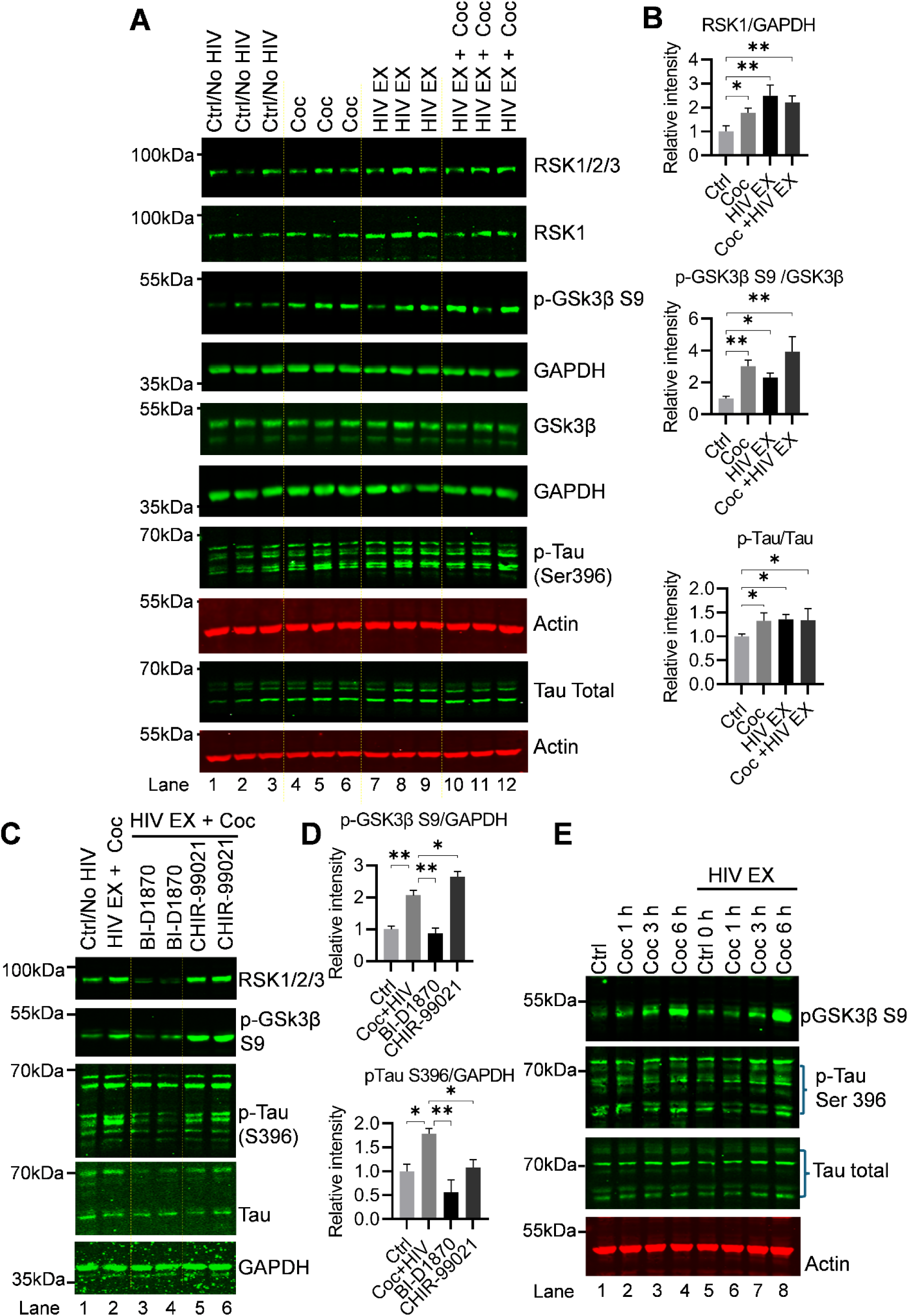
RSK1 functions upstream of GSK3β to mediate Tau phosphorylation induced by HIV exposure and cocaine. (**A & B**) H80 cells were cultured in twelve independent dishes (three biological replicates per condition) under each identical condition and whole-cell lysates were collected 48 h after cocaine, HIV exposure and cocaine plus HIV exposure. Protein lysates were prepared separately from each dish and quantified. Equal amounts of protein were loaded and analyzed by immunoblotting. Immunoblot analysis revealed a significant increase in Tau phosphorylation at Ser396 (p-Tau-S396), a marker of pathological Tau, and RSK-1 expression under all conditions (cocaine, HIV exposure and both) compared with untreated controls. Concurrent analysis demonstrated enhanced phosphorylation of GSK3β at Ser9 (p-GSK3β-Ser9), indicating functional inactivation of GSK3β under all conditions, while GSK3β and Tau levels remained unchanged. Densitometric analysis of immunoblots was performed by normalizing band intensities to β actin or total protein, with values expressed relative to the control (Ctrl/No HIV). (**C & D**) To delineate the relative contributions of RSK1 and GSK3β, H80 cells were pretreated for 24 h with selective inhibitors BI-D1870 (RSK1 inhibitor) or CHIR-99021 (GSK3β inhibitor) prior to HIV and/or cocaine exposure. Immunoblotting and densitometry revealed that BI-D1870 effectively suppressed RSK1 activation and significantly reduced Tau phosphorylation, whereas CHIR-99021 failed to alter Tau phosphorylation induced by HIV or cocaine, confirming that Tau modification is primarily mediated through RSK1-dependent signaling. Inhibition of RSK1 also reversed GSK3β inactivation, as evidenced by reduced Ser9 phosphorylation, suggesting a hierarchical relationship in which RSK1 lies upstream of GSK3β. Densitometric analysis of immunoblots was performed by normalizing band intensities to GAPDH, with values expressed relative to the control (Ctrl/No HIV). **(E)** An acute time point study was conducted for 1 h, 3 h and 6 h with cocaine and cocaine along with HIV, Immunoblot results show enhanced phosphorylation of GSK3β at Ser9 (p-GSK3β-Ser9), indicating functional inactivation of GSK3β at 6h while simultaneously seen the enhanced tau phosphorylation at S396. Immunoblots are representative of at least three independent biological replicates. Data are presented as mean ± S.D. Statistical significance was assessed using one way ANOVA with Dunnett’s multiple comparisons test. Significance is indicated as P < 0.05 (*) and P < 0.01 (**).

Furthermore, to delineate the relative contributions of RSK1 and GSK3β to Tau phosphorylation in response to exposures HIV and cocaine, we employed highly specific small molecular pharmacological inhibitors. Our rationale was that if RSK1 is the common mediator of Tau phosphorylation induced by these stimuli, then its inhibition should suppress this effect, whereas inhibition of GSK3β would not. H80 cells were pretreated for 24 hours with BI-D1870 (a selective RSK1 inhibitor) or CHIR-99021 (a highly specific GSK3β inhibitor), followed by exposure to HIV, cocaine, or both. After treatment, total protein lysates were analyzed by immunoblotting to assess signaling and phosphorylation dynamics (**Figure 6C and 6D**). Consistent with our earlier findings (**Figures 2 and 3**), both HIV and cocaine exposures resulted in robust upregulation of RSK1 activity compared to untreated controls (**Figure 6C and 6D**). Importantly, pretreatment with BI-D1870 effectively suppressed RSK1 activation, whereas CHIR-99021 had no effect on RSK1 activity, indicating that RSK1 functions independently of, and upstream from, GSK3β in this signaling cascade (**Figure 6C and 6D**). We next examined the effect of these inhibitors on GSK3β activity. As shown previously (**Figure 4**), both HIV and cocaine exposures led to inactivation of GSK3β, as evident from enhanced phosphorylation at S9 (**Figure 6C and 6D**). Notably, inhibition of RSK1 with BI-D1870 reduced Ser9 phosphorylation, suggesting restoration of GSK3β activity and indicating that RSK1 contributes to GSK3β inactivation (**Figure 6C and 6D**). In contrast, direct inhibition of GSK3β with CHIR-99021 resulted in sustained inactivation, confirming the specificity and effectiveness of the inhibitors. To evaluate the downstream effects of these perturbations, we examined Tau phosphorylation and found that both HIV and cocaine exposure markedly increased Tau phosphorylation, consistent with RSK1 activation. (**Figures 6C and 6D**).

Importantly, RSK1 inhibition with BI-D1870 markedly suppressed Tau phosphorylation, whereas inhibition of GSK3β with CHIR-99021 did not alter Tau phosphorylation induced by HIV exposure or cocaine. These findings demonstrate that Tau phosphorylation in this context is primarily mediated through an RSK1-dependent, GSK3β-independent mechanism. Interestingly, prolonged treatment (≥24 hours) with BI-D1870 also resulted in a reduction in total RSK1 protein levels, while housekeeping controls (GAPDH) remained unchanged. This suggests that sustained pharmacological inhibition may influence not only RSK1 activity but also its protein stability or turnover.

Collectively, these results establish a hierarchical signaling relationship in which RSK1 acts upstream of GSK3β and plays a central role in mediating Tau phosphorylation following HIV and cocaine exposure. Furthermore, they highlight RSK1 as a critical therapeutic target, as its inhibition effectively attenuates Tau pathology while also modulating downstream kinase signaling. We next investigated the temporal dynamics of acute cocaine- and HIV-induced signaling in H80 cells to determine whether rapid inactivation of GSK3β, reflected by increased phosphorylation at the inhibitory Ser9 site, occurs in parallel with changes in Tau phosphorylation at the pathological Ser396 residue. To assess time-dependent effects, cells were exposed to cocaine or HIV for 1, 3, and 6 hours (**Figure 6E**). Protein lysates collected at each time point were analyzed by immunoblotting for p-GSK3β Ser9, p-Tau Ser396, total Tau, and actin. Both cocaine and HIV induced a sustained increase in GSK3β Ser9 phosphorylation at the 3- and 6-hour time points (**lanes 3–4 vs. lane 1; lanes 7–8 vs. lane 5**), indicating persistent inhibition of GSK3β activity during acute exposure. Notably, this inhibitory modification did not lead to a reduction in Tau phosphorylation. Instead, we observed a progressive and robust increase in p-Tau Ser396 over time, demonstrating that Tau phosphorylation continues to accumulate despite functional inactivation of GSK3β. The simultaneous suppression of GSK3β activity and enhancement of Tau phosphorylation provides strong evidence for a GSK3β-independent mechanism of Tau regulation under both cocaine and HIV exposure. These findings implicate alternative kinases, most prominently RSK1, as key drivers of Tau phosphorylation at Ser396, even in the absence of active GSK3β. Collectively, these data demonstrate that acute cocaine and HIV exposures sustain Tau hyperphosphorylation independently of GSK3β activity, highlighting RSK1 as a dominant upstream kinase in this process. These results are consistent with our chronic exposure studies, further reinforcing a model in which RSK1-dependent signaling persistently drives Tau phosphorylation across temporal contexts.

### RSK1 knockout impairs AKT signaling, activates GSK3β, and suppresses Tau phosphorylation

To further confirm the direct role of RSK1 in regulating Tau phosphorylation, we generated RSK1 knockout (KO) H80 cells using CRISPR-Cas9. Immunoblot analysis confirmed efficient loss/reduction of RSK1 protein, enabling us to investigate the downstream signaling pathways. Strikingly, RSK1 ablation led to a marked reduction in Tau phosphorylation at Ser396 (p-Tau S396), while total Tau levels remained unchanged, indicating that RSK1 regulates Tau primarily through post-translational mechanisms rather than transcriptional or translational control. Quantitative analyses across independent experiments confirmed a significant decrease in p-Tau S396 in RSK1 KO cells compared with controls (**Figure 7A and B**), establishing that RSK1 is required for efficient phosphorylation of Tau at this pathological site.

**Figure 7:**
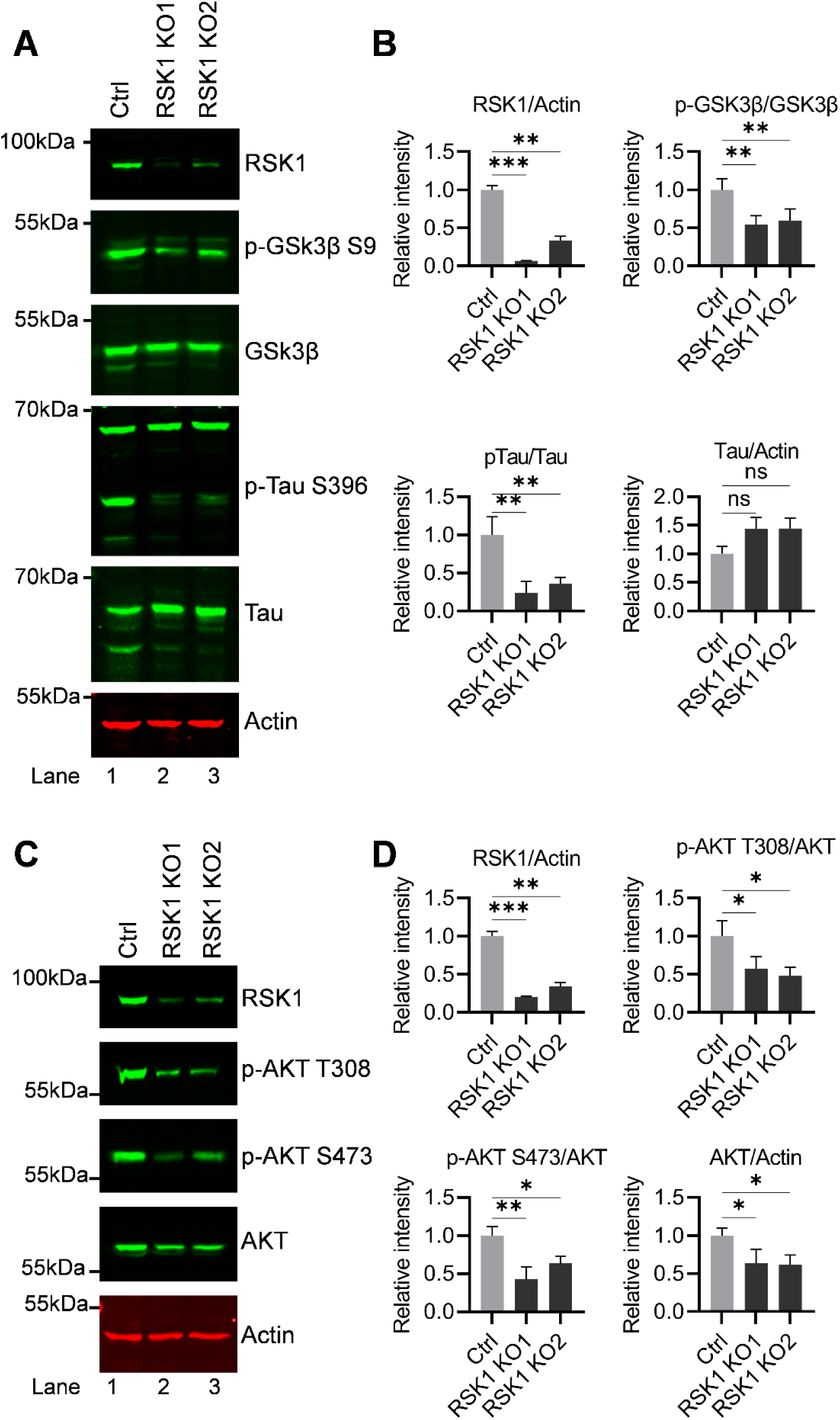
CRISPR-Cas9-mediated RSK1 knockout reveals its essential role in Tau phosphorylation and AKT-GSK3β signaling. / RSK1 as a critical upstream regulator of Tau phosphorylation and AKT signaling, functioning hierarchically above GSK3β and contributing to multiple signaling pathways relevant to neurodegeneration. (**A–B**) Immunoblot analysis confirmed successful RSK1 knockout (RSK1 KO) in H80 cells and demonstrated reduced inhibitory phosphorylation of GSK3β at Ser9 (p-GSK3β-Ser9), indicating reactivation of GSK3β kinase activity upon loss of RSK1. RSK1 KO also led to a significant reduction in Tau phosphorylation at Ser396 (p-Tau-S396) compared with control cells, while total Tau levels remained unchanged. Despite GSK3β reactivation, Tau phosphorylation did not recover, suggesting that RSK1 mediates site-specific Tau phosphorylation independently of GSK3β. Quantitative analysis from multiple independent experiments confirmed a significant decrease in p-Tau-S396 in RSK1 KO cells, establishing RSK1 as essential for efficient Tau phosphorylation. **(C-D)** Analysis of AKT signaling revealed that RSK1 knockout severely impaired AKT activation, as evidenced by a marked reduction in phosphorylation at Thr308 and Ser473, and also decreased total AKT protein levels. These findings indicate that RSK1 positively regulates both AKT activation and AKT protein stability. Densitometric analysis of immunoblots was performed by normalizing band intensities to β actin or total protein, with values expressed relative to the control. Immunoblots are representative of at least three independent biological replicates. Data are presented as mean ± S.D. Statistical significance was assessed using one way ANOVA with Dunnett’s multiple comparisons test. Significance is indicated as P < 0.05 (*), P < 0.01 (**) and P < 0.001 (***).

Interestingly, RSK1 knockout also impacted GSK3β signaling. Specifically, loss of RSK1 resulted in a reduction of inhibitory phosphorylation of GSK3β at Ser9, indicating reactivation of GSK3β kinase activity. These findings demonstrate that RSK1 acts upstream of GSK3β and contributes to its inactivation, consistent with our pharmacological inhibition data (**Figure 6**). Notably, however, reactivation of GSK3β did not restore Tau phosphorylation at Ser396, strongly documenting that RSK1 drives site-specific Tau phosphorylation independently of GSK3β (**Figure 7**). This observation highlights the complexity of Tau regulatory networks and indicates that RSK1 is the main kinase controlling pathological Tau modification (Tau-S396), rather than merely modulating canonical Tau kinases such as GSK3β.

We further evaluate the impact of RSK1 loss on AKT signaling pathway. RSK1 knockout resulted in severely impaired AKT signaling pathway, as evidenced by a marked reduction in phosphorylation at both Thr308 and Ser473. Notably, total AKT protein levels were also reduced, suggesting that RSK1 contributes not only to AKT activation but also to AKT protein stability and/or abundance. These findings identify RSK1 as a positive regulator of AKT signaling at both functional and protein stability levels (**Figure 7C and D**). Collectively, the coordinated effects of RSK1 deletion, including suppression of Tau phosphorylation, reactivation of GSK3β, and attenuation of AKT signaling, establish RSK1 as a central upstream regulator of interconnected kinase networks governing Tau pathology. Importantly, the inability of GSK3β reactivation to rescue Tau phosphorylation further underscores the primary role of RSK1 in mediating Tau-S396 phosphorylation.

Taken together, these results position RSK1 as a critical signaling hub integrating AKT and GSK3β pathways to regulate Tau phosphorylation and identify it as a promising therapeutic target for Tauopathies, including HIV-associated neurocognitive disorders (HAND) and cocaine-associated neurodegeneration.

### RSK1 functions as an upstream regulator of AKT- GSK3β signaling cascade

As detailed above, we identified RSK1 as a key upstream regulator of both AKT and GSK3β signaling in H80 cells. Loss (knock out) or pharmacological inhibition of RSK1 impaired AKT activation and simultaneously reactivated GSK3β, as evidenced by reduced phosphorylation at its inhibitory Ser9 site. To further substantiate this regulatory hierarchy, we next tested whether RSK1 overexpression produces the reciprocal effects. Based on our prior findings, we hypothesized that elevated RSK1 levels would enhance AKT activation while promoting GSK3β inactivation via increased Ser9 phosphorylation.

To evaluate the signaling consequences of RSK1 upregulation, H80 cells were transiently transfected with a CMV-driven RSK1 expression construct, and lysates were collected 48 hours post-transfection for immunoblot analysis. Overexpression was confirmed by a marked increase in total RSK1 protein, along with elevated levels of RSK1/2/3 isoforms, validating robust induction of the RSK signaling axis (**Figure 8A**). Notably, phosphorylation of RSK1 at Ser380, a key autophosphorylation site associated with catalytic activation, was significantly increased. However, phosphorylation at Thr348, a critical activation loop residue in the N-terminal kinase domain that is typically phosphorylated downstream of ERK signaling, remained unchanged or even decreased relative to vector-transfected controls. The selective increase in pSer380 without a corresponding increase in pThr348 suggests that RSK1 overexpression alone does not result in full enzymatic activation and may reflect a partial or ERK-independent activation state. This phosphorylation pattern implies that RSK1 may be primed for signaling but is not fully engaged in substrate phosphorylation, potentially limiting canonical downstream activity.

**Figure 8:**
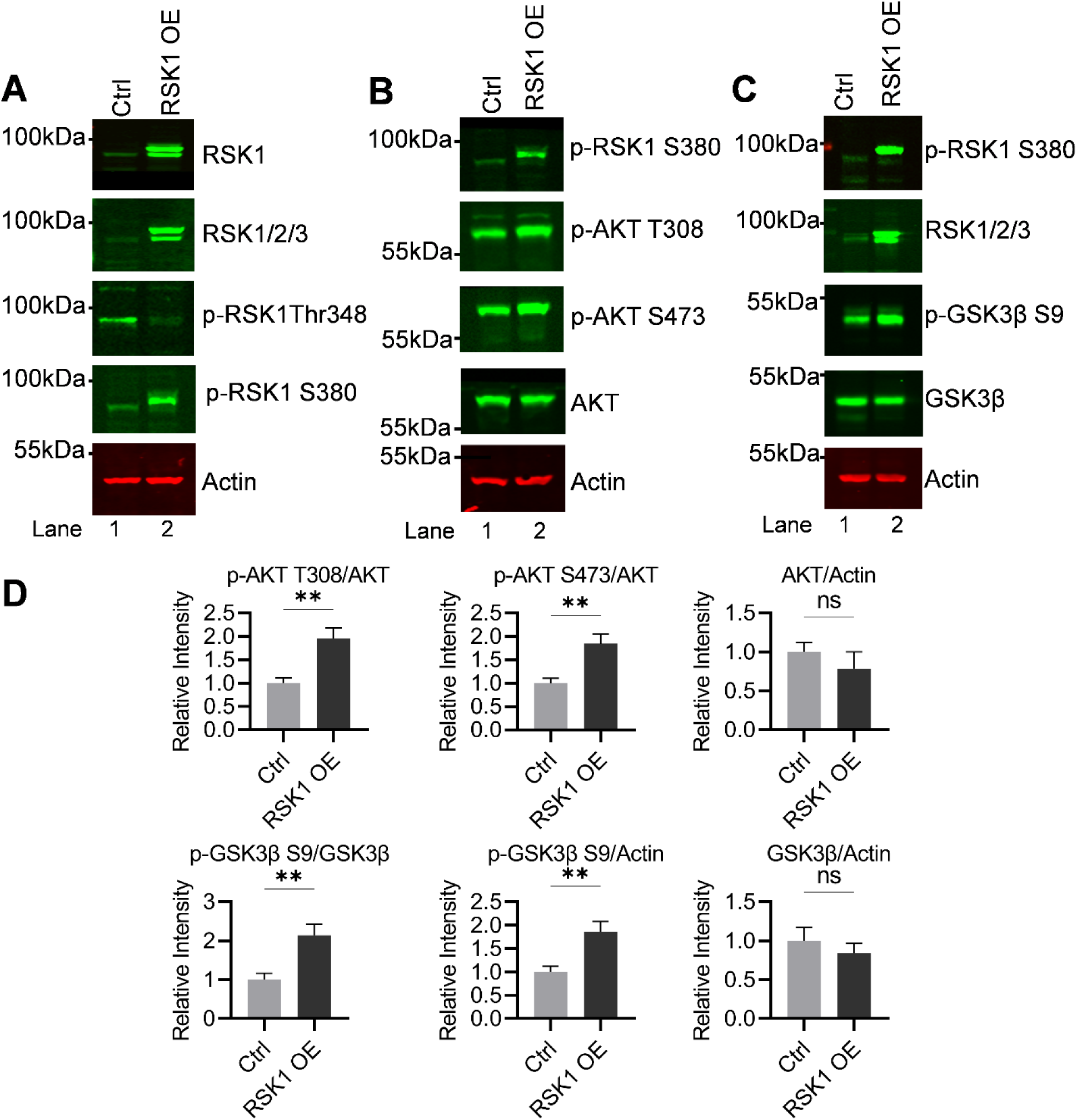
RSK1 overexpression modulates AKT and GSK3β signaling in H80 cells. / RSK1 as a central upstream modulator of AKT and GSK3β signaling. (**A–B**) Immunoblot analysis confirmed robust overexpression of RSK1 following transient transfection with a CMV-driven RSK1 construct, as evidenced by a marked increase in total RSK1 protein and elevated levels of RSK isoforms (RSK1/2/3). Overexpression also resulted in a significant increase in phosphorylation of RSK1 at Ser380, an autophosphorylation site associated with partial catalytic activation, while phosphorylation at Thr348 decreased relative to empty vector transfected control. **(C–D)** RSK1 overexpression induced a robust increase in AKT phosphorylation at both Thr308 and Ser473, two canonical regulatory sites required for full AKT activation, without altering total AKT protein levels. These findings indicate that RSK1 positively regulates AKT activity predominantly through post-translational mechanisms. (**E–F**) Overexpression of RSK1 also enhanced phosphorylation of GSK3β at Ser9, a canonical inhibitory site that suppresses GSK3β kinase activity, while total GSK3β levels remained unchanged. Densitometric analysis of immunoblots was performed by normalizing band intensities to β actin or total protein, with values expressed relative to the empty vector transfected control. Immunoblots are representative of at least three independent biological replicates. Data are presented as mean ± S.D. Statistical significance was assessed using an unpaired, two tailed Student’s t test. Significance is indicated as P < 0.01 (**) and ns denotes not significant.

Consistent with our model, RSK1 overexpression resulted in robust activation of AKT, as demonstrated by increased phosphorylation at both Thr308 and Ser473, the two critical regulatory sites required for full AKT activity (**Figure 8B and D**). Importantly, total AKT protein levels remained unchanged, indicating that RSK1 enhances AKT signaling primarily through post-translational activation rather than changes in protein abundance.

In parallel, RSK1 overexpression led to a pronounced increase in inhibitory phosphorylation of GSK3β at Ser9 (**Figure 8C and D**), confirming functional inactivation of this kinase. Total GSK3β levels remained constant, further supporting that this effect reflects post-translational regulation. These findings reinforce the conclusion that RSK1 negatively regulates GSK3β activity through phosphorylation-dependent inhibition, likely in part via AKT activation.

Collectively, these results establish RSK1 as a central upstream modulator of the AKT-GSK3β signaling axis in H80 cells. By simultaneously activating AKT and suppressing GSK3β, RSK1 integrates MAPK/RSK and PI3K/AKT signaling pathways and creates a cellular environment conducive to pathological Tau phosphorylation. This mechanistic link suggests that dysregulation of RSK1 could shift neuronal kinase networks toward pathological Tau modification, with broader implications for survival, metabolism, and neurodegenerative processes. This coordinated regulation highlights RSK1 as a critical signaling hub that governs kinase network balance and promotes Tau dysregulation. Altogether, our data position RSK1 as a key mechanistic driver and potential therapeutic target in conditions characterized by aberrant Tau phosphorylation, including HIV-mediated neurotoxicity (HAND) and cocaine-induced neurodegeneration.

### Cocaine- and HIV-induced signaling is conserved across 2D neuronal cultures, 3D spheroids, and brain organoid model systems

To determine whether the signaling pathways induced by cocaine and HIV in H80 cells are conserved across additional neuronal systems, we extended our investigation to SH-SY5Y neuroblastoma cells (**Figure 9B**), 3D neuronal spheroids (**Figure 9A and 9C**), and human iPSC-derived brain organoids (**Figure 9D**). This approach allowed us to evaluate the robustness and reproducibility of the identified signaling axis across models of increasing biological complexity.

**Figure 9:**
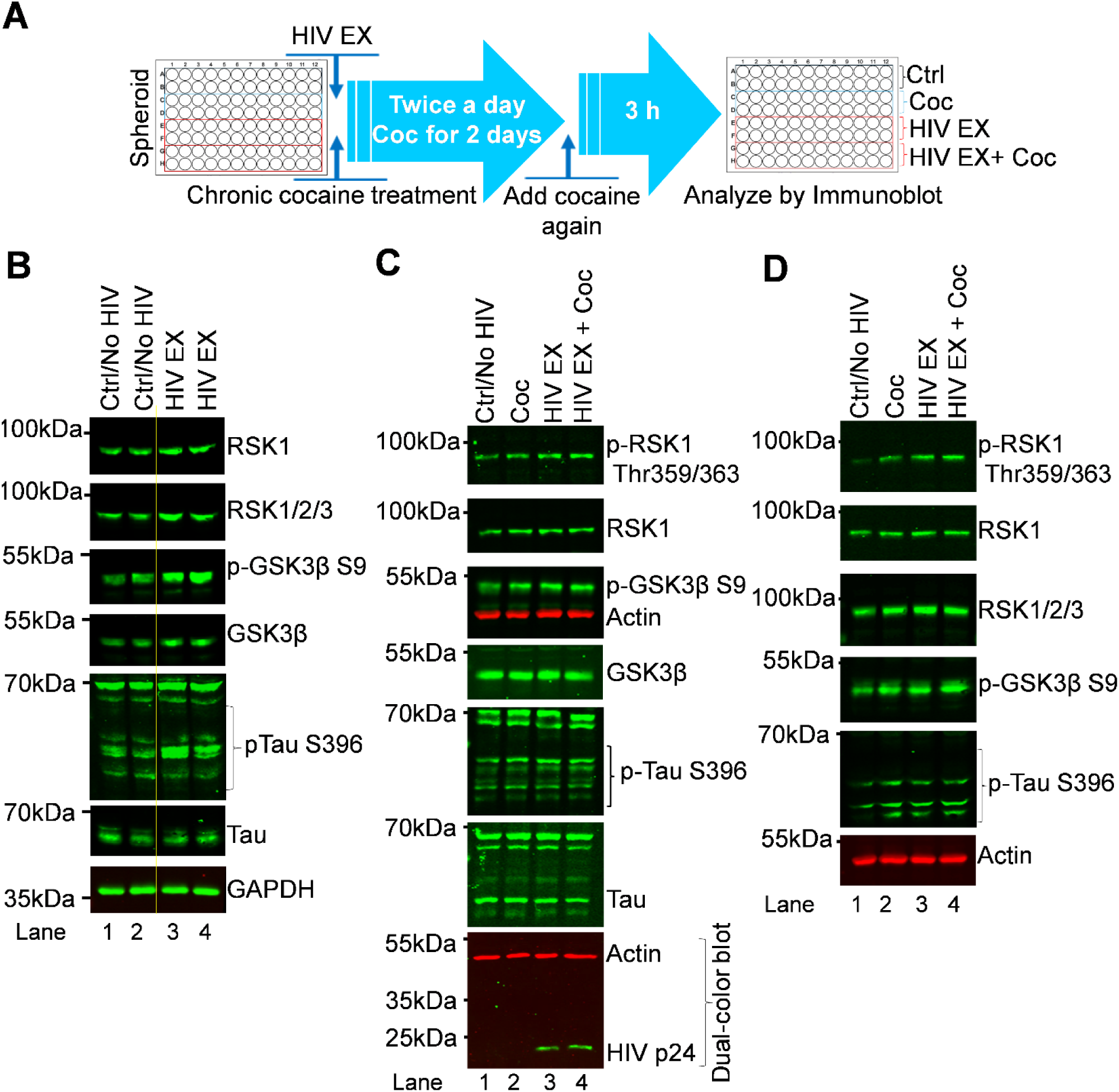
HIV exposure and cocaine induce RSK1–dependent Tau phosphorylation in neuronal monolayers, 3D spheroid, and brain organoid models. (**A**) Schematic representation of the immunoblot experimental workflow illustrating HIV and cocaine exposure in a three–dimensional spheroid culture system. (**B**) SH–SY5Y neuronal cells were exposed to HIV for 48 h, followed by cell lysis and immunoblot analysis. HIV exposure resulted in upregulation of RSK1, increased inhibitory phosphorylation of GSK3β at Ser9, and enhanced phosphorylation of Tau at Ser396. **(C)** These immunoblot findings were recapitulated in a 3D multicellular spheroid model composed of H80 neurons, microglia, and SH–SY5Y cells, confirming the reproducibility of HIV–induced signaling responses in a heterogeneous cellular context. **(D)** Immunoblot analysis in a three–dimensional organoid model further validated HIV– and cocaine–induced activation and regulation of RSK1, phosphorylation GSK3β S9, and Tau phosphorylation S396, demonstrating the robustness of this signaling axis across increasingly complex neuronal systems.

First, to assess reproducibility in an independent neuronal cell line, SH-SY5Y cells were exposed to HIV for 48 hours, followed by immunoblot analysis of key signaling markers. Chronic HIV exposure resulted in a pronounced upregulation of RSK1, accompanied by increased inhibitory phosphorylation of GSK3β at Ser9, while total GSK3β levels remained unchanged (**Figure 9B; lanes 3–4 vs. 1–2**). Notably, this was paralleled by a significant increase in Tau phosphorylation at Ser396, demonstrating that HIV induces a coordinated signaling response involving RSK1 activation, GSK3β inactivation, and Tau dysregulation. These findings confirm that the signaling axis identified in H80 cells is reproducible in additional neuronal cell types.

To further examine whether these mechanisms are preserved in a more physiologically relevant 3D context, we utilized a multicellular neuronal spheroid model. Spheroids were generated by co-culturing equal numbers of H80 cells, microglia, and SH-SY5Y cells (15,000 cells total per spheroid), and subjected to control, cocaine, HIV, or combined treatments. Following 48-hour exposure, pooled spheroids from each condition were analyzed by immunoblotting. Consistent with 2D models, both cocaine and HIV treatments induced robust RSK1 upregulation, increased GSK3β Ser9 phosphorylation, and enhanced Tau phosphorylation at Ser396 (**Figure 9C**). Importantly, the presence of microglia enabled productive HIV infection within the spheroid system, further increasing physiological relevance. These coordinated molecular changes demonstrate that the RSK1-GSK3β-Tau signaling axis is preserved within a multicellular 3D neuronal microenvironment.

We next evaluated whether these findings extend to higher-order neural systems using human cerebral organoids (hCOs) derived from hiPSCs. Following exposure to cocaine, HIV, or both, organoids were processed for immunoblot analysis. Consistent with results from both 2D cultures and spheroids, treated organoids exhibited marked upregulation of RSK1, increased inhibitory phosphorylation of GSK3β at Ser9, and sustained Tau phosphorylation at Ser396 (**Figure 9D**). Notably, Tau phosphorylation remained elevated despite GSK3β inactivation, reinforcing the presence of a GSK3β-independent mechanism, likely mediated by RSK1. These results confirm that the identified signaling pathway is conserved even in complex, human-relevant 3D brain models.

Together, these data demonstrate that the core signaling cascade, RSK1 activation/upregulation, GSK3β inactivation, and pathological Tau phosphorylation, identified in H80 cells is highly reproducible across neuronal systems of increasing complexity, including 3D cultures, multicellular spheroids, and brain organoids. This consistency underscores the biological robustness and generalizability of this pathway and highlights its relevance across diverse human-derived neural contexts, strengthening its potential significance in HIV- and cocaine-associated neurodegeneration.

Overall, our investigation identifies RSK1 as a central signaling hub that integrates and coordinates multiple kinase pathways governing Tau phosphorylation. Both HIV and cocaine robustly activate RSK1, which eventually promotes the inactivation of GSK3β, establishing a convergent downstream signaling axis despite distinct upstream regulatory mechanisms. Importantly, Tau phosphorylation persists even under conditions of GSK3β inhibition, demonstrating that RSK1 drives pathological Tau modification through a GSK3β-independent mechanism. These findings establish RSK1 as an essential upstream regulator of interconnected kinase networks that control site-specific Tau phosphorylation.

Notably, these signaling dynamics are consistently reproduced across multiple neuronal systems, including 2D cultures, 3D spheroids, and human brain organoids, underscoring the robustness, reproducibility, and biological relevance of this pathway across diverse neural contexts. Collectively, these results support a unified model in which RSK1 serves as the primary mediator linking HIV and cocaine exposure to Tau dysregulation and neuronal stress responses. Beyond HAND and substance use-related neurotoxicity, this mechanism has broader implications for Tau-driven neurodegenerative diseases, including Alzheimer’s disease and related cognitive disorders. Thus, RSK1 emerges as a key mechanistic driver and a promising therapeutic target for conditions characterized by aberrant Tau phosphorylation and neurodegeneration.

### Distinct yet convergent signaling mechanisms by which HIV exposure and cocaine drive Tau phosphorylation/ pathology

To summarize our findings, we propose the following model for HIV- and cocaine-induced Tau phosphorylation (**Figure 10**). In this study, we demonstrate that exposure to HIV and cocaine leads to robust Tau phosphorylation, and we delineate the distinct yet convergent molecular mechanisms underlying this process. Although both stimuli ultimately induce Tau phosphorylation, the upstream signaling pathways they engage are mechanistically distinct. In the context of HIV exposure, we observed a pronounced and sustained upregulation and activation of RSK1. Activated RSK1 promotes Tau phosphorylation while simultaneously inhibiting GSK3β activity through an AKT-independent mechanism. Consistent with this pathway, HIV exposure resulted in a marked increase in Tau phosphorylation, identifying RSK1 as a dominant mediator of HIV-driven Tau dysregulation. On the other hand, cocaine exposure engages a partially overlapping but distinct signaling cascade. While cocaine induces modest activation of RSK1, it strongly stimulates AKT, as evidenced by robust phosphorylation at Thr308 and Ser473, both required for full catalytic activation. Activated AKT subsequently phosphorylates GSK3β at Ser9 (p-GSK3β-Ser9), leading to its functional inactivation. Despite differences in upstream signaling intensity, cocaine also promotes Tau phosphorylation, highlighting a mechanism that does not rely solely on the robust RSK1 activation observed in the case of HIV. Notably, Tau phosphorylation persists even under conditions of GSK3β inactivation in both HIV- and cocaine-exposed systems. This finding reveals the existence of a GSK3β-independent mechanism of Tau modification and establishes RSK1 as a key regulator of Tau phosphorylation under these conditions. The persistence of Tau phosphorylation despite suppression of canonical GSK3β activity suggests the involvement of parallel or compensatory signaling pathways that warrant further investigation.

**Figure 10:**
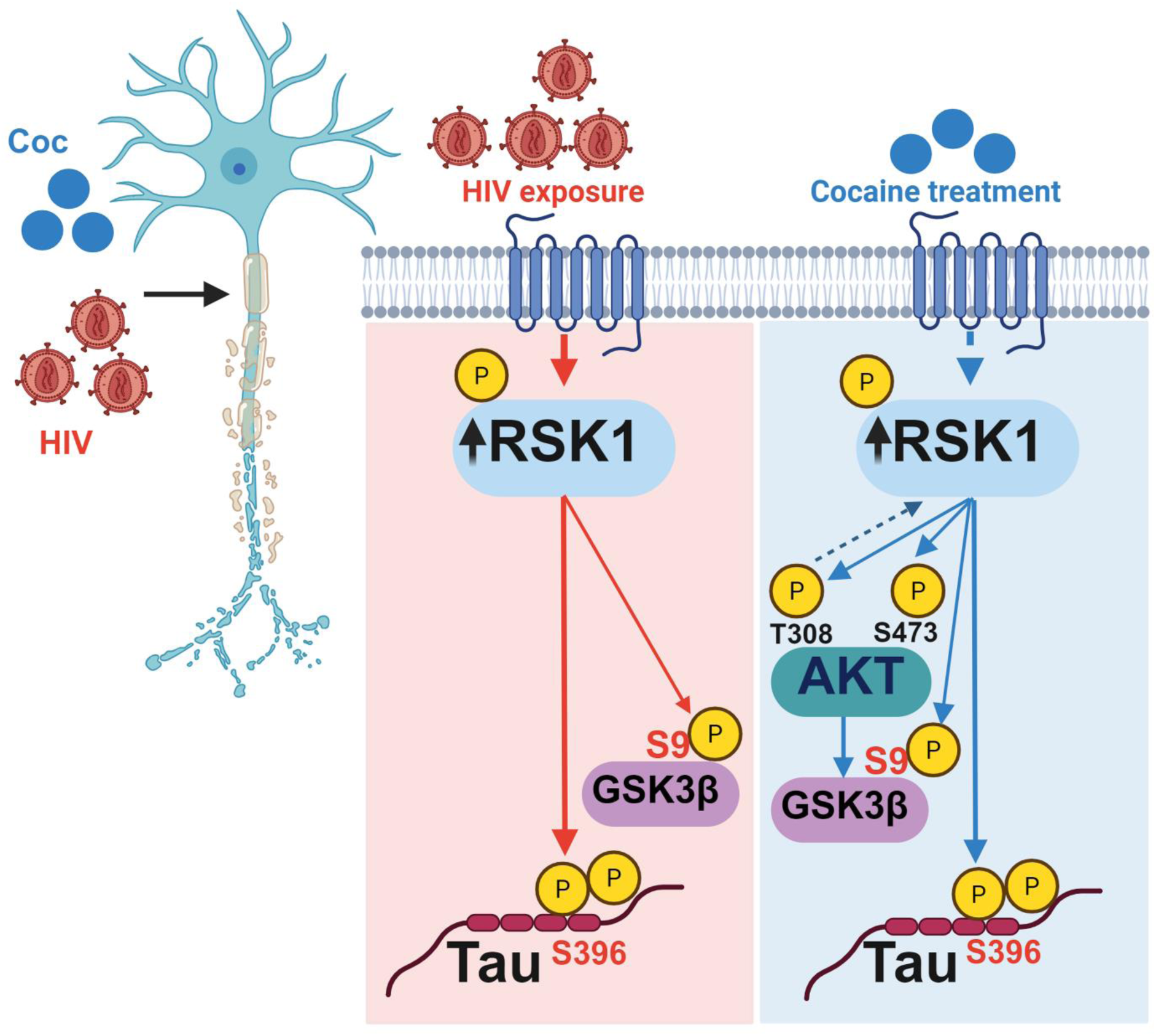
Model summarizing HIV– and cocaine–induced Tau phosphorylation in neuronal cells. Proposed schematic illustrating distinct yet convergent signaling mechanisms by which HIV exposure and cocaine promote Tau phosphorylation in H80 neuronal cells. HIV exposure induces a robust and sustained activation and upregulation of RSK1, which drives Tau phosphorylation through a pathway that is independent of AKT signaling while concurrently promoting inhibitory phosphorylation of GSK3β at Ser9. In contrast, cocaine exposure engages a partially overlapping but mechanistically distinct pathway, characterized by modest RSK1 induction and strong activation of AKT, as evidenced by phosphorylation at Thr308 and Ser473. Activated AKT subsequently catalyzes inhibitory phosphorylation of GSK3β at Ser9, leading to its functional inactivation. Despite GSK3β inactivation under both conditions, Tau phosphorylation persists, indicating the existence of a GSK3β–independent mechanism regulated by RSK1. Collectively, these findings identify RSK1 as a central signaling hub that integrates viral and substance–induced signaling to drive Tau dysregulation, highlighting its critical role in neurodegenerative processes relevant to HIV–associated neurocognitive disorders.

Importantly, we identify RSK1 as a critical upstream regulator of both AKT and GSK3β signaling, exerting positive control over AKT activation while negatively regulating GSK3β activity. This dual regulatory capacity positions RSK1 as a central signaling hub integrating viral and drug-induced pathways that converge on Tau pathology. Collectively, our findings provide mechanistic insight into how HIV and cocaine exposure, through distinct yet convergent pathways, drive Tau dysregulation and contribute to neurotoxicity. From a therapeutic perspective, these results highlight RSK1 as a promising target for intervention, offering a unifying framework for mitigating tauopathy in neuroHIV, HAND, and cocaine-associated neurodegeneration.

## Discussion

Neuronal cell lines such as SH–SY5Y are widely used experimental models, yet they have limitations that can compromise experimental robustness [56]. Neurons cells are highly sensitive to culture conditions, exhibit variable growth and differentiation rates, and frequently display batch–to–batch and passage–dependent heterogeneity [64]. Such instability poses significant challenges for studies requiring long–term culture or consistent phenotypic behavior, particularly investigations focused on neurodegeneration, Tau–related pathology, or kinase–driven signaling mechanisms. These constraints highlight the need for neuronal models that retain essential neuron–like properties while offering greater stability, reproducibility, and ease of maintenance. More robust and tractable cell systems not only improve experimental consistency but also enable higher–throughput analyses and more reliable interpretation of signaling mechanisms relevant to neuropathies, tauopathies, and neurotoxic exposures.

In this study, we characterized H80 cells as a stable and experimentally tractable neuronal model and employed them to investigate the molecular mechanisms by which HIV and cocaine promote Tau phosphorylation and neurotoxicity. H80 cells were selected based on their robust proliferation, low cytotoxicity, and stable culture performance. Through immunofluorescence, qPCR, and Western blot analyses, we confirmed that H80 cells express key neuronal markers, including NeuN, MAP2, and Tau, consistent with a mature neuronal phenotype (**Figure 1A-C; Supplementary Figure S1**). Notably, the expression of MAP2, an axon-associated protein critical for neuronal architecture and implicated in neurodegenerative processes, further supports the neuronal identity of H80 cells. Collectively, the presence of these well-established neuronal markers demonstrates that H80 cells possess essential neuron-like features and extends their utility beyond glioma research. Importantly, given our focus on HIV-induced neurotoxicity, we assessed whether H80 cells express key HIV receptors and co-receptors. Our results show that H80 cells do not express CD4 or CCR5, the canonical receptor and major co-receptor for HIV entry, respectively. However, H80 does express one of the HIV co-receptors, CXCR4, in approximately 20% of cells (**Figure 1D**). This expression profile is consistent with previous reports indicating that neurons lack CD4 but express chemokine receptors such as CXCR4 and CCR5. Prior studies, including those by Kaul and colleagues, have demonstrated that although due to the absence of HIV receptor, neurons are not productively infected by HIV, they remain highly susceptible to HIV-induced injury mediated through chemokine receptors and downstream signaling cascade, with CXCR4 serving as a key mediator of neurotoxic signaling in neurons [17, 57]. The absence of CCR5 and the selective expression of CXCR4 in H80 cells therefore provide a unique and focused system to investigate CXCR4-dependent mechanisms of HIV-associated neuronal stress. This receptor profile enables us to dissect how HIV exposure perturbs neuronal signaling in the absence of CCR5-mediated protective pathways, thereby facilitating a clearer understanding of HIV-induced neurotoxicity.

Our study identifies a previously unrecognized mechanistic link between viral exposure and neuronal stress pathways. Using an integrated approach combining transcriptional analysis, immunofluorescence, and biochemical analyses, we demonstrate that exposure to HIV virions robustly activates inflammatory signaling cascades and induces neurotoxic responses in H80 neuronal cells. Specifically, HIV exposure significantly upregulates the transcripts of pro-inflammatory cytokines IL-1β and TNF-α (**Figure 2B**), revealing a previously underappreciated neuron-intrinsic inflammatory response. These findings indicate that direct interaction with viral particles is sufficient to trigger canonical neuroinflammatory programs in neuronal cells. Given that IL-1β and TNF-α are well-established mediators of neuronal injury in both HIV-associated neurocognitive disorders (HAND) and Alzheimer’s disease, our results highlight a shared inflammatory axis between virally induced and classical neurodegenerative processes, characterized by proinflammatory cytokines and pathologic Tau phosphorylation. Although prior studies have primarily attributed HIV-induced cytokine production to microglia [65, 66], emerging evidence suggests that neurons can also produce cytokines that modulate synaptic function and central nervous system homeostasis, a fact further strengthened using our novel neuronal model system. [67, 68]. The robust cytokine induction observed here further supports effective exposure of H80 cells to HIV virions and underscores the capacity of neurons to directly engage in inflammatory signaling. Importantly, our biochemical analyses reveal that HIV-exposed H80 cells exhibit concurrent increases in RSK1 protein levels and Tau phosphorylation at Ser396 (p-Tau S396), with quantitative immunoblotting demonstrating a strong correlation between these events (**Figures 2C-E**). In contrast, total Tau protein levels were only modestly altered, indicating that HIV primarily drives post-translational modification of Tau rather than increasing its overall expression and abundance. These findings support a model in which HIV-induced RSK1 activation directly promotes pathogenic Tau phosphorylation. Given that phosphorylation at Ser396 is closely associated with Tau protein aggregation and neuronal toxicity, our data establishes RSK1 as a critical effector linking HIV exposure to early tauopathic signaling. The simultaneous induction of inflammatory cytokines and RSK1 further suggests coordinated activation of inflammatory and stress-responsive kinase pathways, creating a signaling environment that may initially be adaptive but ultimately becomes maladaptive. As a downstream effector of MAPK signaling, RSK1 plays key roles in transcriptional regulation and cellular stress responses; however, its sustained activation appears to drive pathological Tau modification. Collectively, these findings identify RSK1 as a central molecular node connecting HIV-induced inflammatory signaling to Tau pathology. By delineating this pathway, our study provides new mechanistic insight into how HIV exposure can initiate and accelerate neurodegenerative processes within the central nervous system.

In addition to the effects induced by HIV, our data demonstrate that cocaine also independently activates the RSK1 signaling cascade to drive Tau phosphorylation, revealing a shared, previously unappreciated kinase dependency underlying both viral- and drug-induced neurotoxicity underscoring the convergence of drugs and HIV–mediated stress responses on a shared kinase pathway. Chronic cocaine exposure triggered robust phosphorylation of RSK1 at key regulatory residues (Thr348, Thr359/Ser363, and Ser380), accompanied by a pronounced increase in Tau phosphorylation at Ser396, without any significant change in total Tau protein levels (**Figures 3B–D**). These findings establish that cocaine drives Tau pathology primarily through post-translational mechanisms rather than altering Tau expression via transcriptional or translational regulation. Strikingly, RSK1 activation was both rapid and highly sensitive; even a brief 15-minute exposure to cocaine was sufficient to induce multi-site phosphorylation (**Figures 3E–G**). A comparable activation profile was observed following acute HIV exposure, positioning RSK1 as an immediate and convergent sensor of diverse neurotoxic stimuli. Functional interrogation of RSK1 in mediating Tau phosphorylation unequivocally establishes RSK1 as essential for Tau phosphorylation. Both pharmacological inhibition and genetic ablation of RSK1 completely abolished Tau-Ser396 phosphorylation induced by either HIV or cocaine (**Figure 6**), demonstrating that RSK1 is not merely correlative but a required driver of this process. Notably, these findings add a new layer to the prevailing paradigm that GSK3β is the dominant Tau kinase, instead identifying RSK1 as the principal effector of Tau phosphorylation under conditions of HIV and cocaine exposure. Despite distinct upstream dynamics, HIV and cocaine converge on a common downstream outcome, pathological Tau phosphorylation, through a shared RSK1-centered signaling axis. HIV elicited more robust RSK1 activation than cocaine; however, both stimuli produced comparable levels of Tau phosphorylation, indicating that RSK1 activity, rather than upstream signal intensity, dictates the pathological output. Importantly, combined HIV and cocaine exposure failed to produce additive or synergistic effects, instead reaching a plateau consistent with saturation of a shared signaling pathway. Collectively, these findings redefine the molecular framework of Tau dysregulation by establishing RSK1 as a central and dominant integrator of viral and drug-induced neuronal stress. By orchestrating Tau phosphorylation, RSK1 provides a common node through which diverse upstream perturbations converge on a common pathological outcome. The rapid activation of RSK1 following acute exposure further suggests that it may function as an early sensor of neuronal stress, initiating downstream signaling cascades that culminate in Tau pathology. The rapid activation kinetics further suggest that RSK1 functions as an early molecular sentinel that initiates downstream tauopathic cascades. The convergence of HIV and cocaine on this shared node provides a mechanistic explanation for the heightened vulnerability to neurodegeneration observed in individuals exposed to either insult, particularly in the context of neuroHIV, where comorbid stimulant use is widespread and associated with accelerated cognitive decline. The identification of RSK1 as a unifying mechanistic driver provides a compelling framework for understanding how these interactions may arise and highlights a promising therapeutic target for mitigating Tau pathology across diverse neurotoxic contexts.

An unexpected and conceptually important observation emerged from our investigation, an independent exposure to either HIV or cocaine consistently resulted in inactivation of GSK3β (an increase in phosphorylation at S9), a kinase implicated in driving Tau phosphorylation and neurodegenerative pathology.

This suppression of GSK3β activity was evidenced by a robust increase in its inhibitory phosphorylation at Ser9, yet Tau remained persistently hyperphosphorylated under these same conditions (**Figure 4 and Figure 6**). This strongly suggests that, Tau phosphorylation proceeds through GSK3β–independent mechanisms, thereby elevating, RSK-1. Our findings highlight RSK1 as the dominant kinase responsible for maintaining Tau phosphorylation when GSK3β is rendered inactive. Our data reveal a remarkably consistent pattern, both HIV and cocaine suppress GSK3β activity while Tau phosphorylation remains elevated (**Figure 4 and Figure 6**). This divergence between upstream signaling and Tau modification further supports a model in which RSK1 activation becomes the primary driver of Tau–Ser396 phosphorylation under HIV and cocaine exposure. Importantly, this signaling paradigm proved highly reproducible across multiple experimental conditions. We observed the same pattern of GSK3β inactivation coupled with persistent Tau phosphorylation during acute (**Figure 4C, 4D, 4E, 4F and 6E**) as well as chronic cocaine and HIV exposure (**Figure 4G, 4H, 6A, 6B, 6C and 6D**). HIV and cocaine suppress GSK3β and act through RSK1 to drive Tau phosphorylation, positioning RSK1 as a central mediator of their shared pathogenic effects. Altogether, our findings demonstrate that both cocaine and HIV decrease GSK3β activity and instead engage RSK1 to drive Tau phosphorylation. These results identify RSK1 as a central signaling node through which cocaine and HIV converge to promote shared pathogenic mechanisms.

Further mechanistic dissection revealed a clear divergence in how cocaine and HIV regulate upstream kinase signaling. Cocaine, but not HIV, robustly activated the AKT pathway, as evidenced by increased phosphorylation at Thr308 and Ser473, two critical residues required for full catalytic activation (**Figure 5**). This activation was accompanied by a corresponding increase in inhibitory phosphorylation of GSK3β at Ser9, confirming that cocaine suppresses GSK3β through a canonical AKT-dependent pathway. In contrast, HIV exposure failed to induce measurable AKT activation, indicating that HIV-mediated inhibition of GSK3β proceeds via an AKT-independent mechanism. Instead, HIV selectively upregulates and activates RSK1, which in turn drives GSK3β inactivation. Thus, while both stimuli converge on GSK3β inactivation and Tau hyperphosphorylation, they do so through distinct upstream routes, cocaine engaging both AKT and RSK1, and HIV relying predominantly on RSK1. These findings support a bifurcated signaling model in which cocaine and HIV converge on a shared downstream outcome, GSK3β inhibition and Tau hyperphosphorylation, yet reach this endpoint through mechanistically distinct routes. Cocaine engages both AKT and RSK1, thereby broadly amplifying kinase signaling networks, whereas HIV bypasses AKT entirely and relies predominantly on RSK1 activation. This distinction provides important biological insight: cocaine simultaneously activates survival-associated (AKT) and stress-responsive (RSK1) pathways, while HIV exerts a more targeted yet potent effect through selective RSK1 induction.

Pharmacological and genetic perturbation studies further establish RSK1 as the central regulator of this signaling architecture. Inhibition of RSK1 with BI-D1870 markedly reduced Tau-Ser396 phosphorylation despite restoration of GSK3β activity, demonstrating that RSK1, not GSK3β, is the primary kinase sustaining Tau phosphorylation under both cocaine and HIV exposure (**Figure 6**). Conversely, inhibition of GSK3β with CHIR-99021 had no effect on RSK1 activation, confirming that RSK1 operates upstream of GSK3β in this hierarchy. These findings were further validated by CRISPR-Cas9-mediated knockout of RSK1, which abolished Tau phosphorylation, reactivated GSK3β, and reduced both AKT phosphorylation and total AKT levels (**Figure 7**). Notably, the reduction in AKT abundance following RSK1 depletion suggests that RSK1 contributes to AKT stabilization and activation, placing it at the apex of a coordinated kinase network. Collectively, these results define a unified signaling paradigm in which RSK1 functions as a central integrator linking HIV and cocaine exposure to pathological Tau hyperphosphorylation. This RSK1-driven mechanism operates independently of GSK3β and, in the case of cocaine, is further reinforced by AKT activation, thereby integrating distinct upstream perturbations into a common pathological outcome. The convergence of viral and drug-induced signaling on RSK1 provides a mechanistic explanation for the heightened vulnerability to Tau-associated neurodegeneration observed in neuroHIV, particularly in the context of stimulant drug use.

Importantly, the demonstration that pharmacological inhibition of RSK1 reverses Tau hyperphosphorylation and restores kinase balance underscores its translational potential. These findings position RSK1 as a tractable and previously underappreciated therapeutic target for mitigating Tauopathy in HIV-associated neurocognitive disorders (HAND), as well as in broader neurodegenerative conditions such as Alzheimer’s disease. Future studies in preclinical models, including humanized mouse models, will be essential to determine whether targeting RSK1 can attenuate neuroinflammation, prevent Tau pathology, and ultimately slow or halt neurodegenerative progression.

Prior studies have shown that the Tat protein of HIV can activate GSK3β and contributes to Tat-mediated neurotoxicity [19, 69, 70]. However, in this study we examined neuronal responses to intact, replication-competent HIV virions (HIV-1 strain 93/TH/051; dual-tropic, R5/X4), thereby capturing the integrated effects of the full viral particle without infection and replication. Notably, both acute and chronic exposure to these dual-tropic virions consistently resulted in functional inactivation of GSK3β, as evidenced by increased phosphorylation at the inhibitory Ser9 site (**Figures 4 and 6**). These observations likely reflect the complex composition of intact virions, which contain multiple structural and accessory proteins capable of exerting both activating and inhibitory influences on intracellular kinase networks, ultimately shifting the balance toward GSK3β suppression. Strikingly, cocaine exposure produced a similar biochemical signature, robust Ser9 phosphorylation and inactivation of GSK3β. This convergence suggests that HIV and cocaine, despite engaging distinct upstream signaling pathways, AKT-dependent in the case of cocaine and AKT-independent for HIV, ultimately suppress GSK3β through a shared downstream mechanism. Our data identify RSK1 as the central integrator of this process, coordinating GSK3β inhibition and sustaining Tau phosphorylation. These findings indicate that virion-mediated effects are not merely additive but instead converge on a common RSK1-driven intracellular signaling axis that governs neuronal stress responses and Tau pathology.

Future studies will focus on defining the specific viral determinants within the intact virion that initiate RSK1 activation, as well as identifying the neuronal receptors involved, including the potential role of CXCR4-mediated signaling. Elucidating the precise regulatory sites on RSK1 that mediate downstream suppression of GSK3β, particularly those governing Ser9-directed inhibitory phosphorylation, while simultaneously sustaining Tau phosphorylation at Ser396 will be critical for resolving the hierarchical organization of this signaling network. In parallel, comparative analyses of cocaine- and HIV-mediated pathways will be essential to determine whether these distinct stimuli converge on shared upstream sensors or utilize overlapping signaling modules. Such investigations will help establish whether a common molecular node integrates viral and environmental stressors to modulate neuronal vulnerability.

Importantly, our findings identify RSK1 as a major effector of HIV-induced Tau phosphorylation at Ser396, establishing a direct mechanistic link between viral exposure and tauopathic processes (**Figure 2**). HIV exposure activates inflammatory signaling cascades, leading to upregulation of RSK1 and subsequent phosphorylation of Tau at Ser396, a modification strongly associated with neurofibrillary tangle formation in Alzheimer’s disease and HAND. The concurrent induction of pro-inflammatory cytokines, including IL-1β and TNF-α, further implicates inflammatory stress as a key upstream driver of this pathway. Collectively, these results support a model in which HIV-induced inflammatory and stress-responsive signaling converge on RSK1 to drive Tau pathology. This RSK1-centered mechanism provides a unifying framework linking viral exposure, neuroinflammation, and neurodegeneration, and offers important insight into how HIV infection may initiate or accelerate Tau-mediated neuronal dysfunction within the CNS.

In summary, our findings support a unified signaling network in which RSK1 functions as a central molecular node linking both HIV and cocaine exposure to Tau hyperphosphorylation. We show that Tau phosphorylation can be sustained through a GSK3β independent mechanism under conditions of HIV and cocaine induced stress, extending current models of Tau regulation. In the context of cocaine exposure, our data indicates the presence of a dual axis signaling architecture in which RSK1 cooperates with AKT, integrating stress responsive and survival signaling pathways into a convergent downstream outcome. Collectively, these results indicate that RSK1 is not merely associated with Tau dysregulation but plays a functional role in mediating Tau phosphorylation in the setting of HIV and cocaine exposure.

Through complementary genetic and pharmacological approaches, we demonstrate that RSK1 acts as a dominant and convergent driver of Tau pathology, redefining the hierarchical organization of kinase signaling networks that govern Tau modification. These findings fill a critical gap in our understanding of how diverse upstream insults, viral infection, and substance abuse, converge on shared molecular pathways to drive neurodegeneration. The significance of this work lies in its ability to bridge mechanistic and clinical observations. HIV infection and stimulant use are independently associated with accelerated cognitive decline, yet the molecular basis of their interaction has remained poorly understood. By identifying RSK1 as a unifying signaling hub, our study provides a mechanistic framework that explains how these factors synergistically promote Tauopathy in HIV-associated neurocognitive disorders (HAND) and related neurodegenerative conditions, including Alzheimer’s disease (AD). This conceptual advance establishes a new foundation for investigating the intersection of neuroHIV and substance abuse–associated neuropathology.

Importantly, our findings have immediate translational implications. We demonstrate that pharmacological inhibition of RSK1 reverses Tau hyperphosphorylation and restores kinase homeostasis, identifying RSK1 as a tractable and high-value therapeutic target. Targeting RSK1 offers the potential to intercept pathogenic signaling cascades upstream of irreversible neuronal damage, representing a fundamentally new strategy for mitigating Tau-driven neurodegeneration. Future studies will focus on evaluating the therapeutic efficacy of RSK1 inhibition in physiologically relevant preclinical models, including humanized mouse systems, to determine whether targeting this pathway can attenuate neuroinflammation, suppress Tau pathology, and preserve neuronal function. Successful validation of this approach has the potential to transform therapeutic strategies for HAND and other Tau-associated disorders by targeting a shared and central molecular driver of disease.

## Limitation

The main limitation of the study is that while NeuN, MAP2, and Tau serve as well-established neuronal markers, future studies should incorporate additional proteins associated with synaptic activity and neuronal function, such as synaptophysin, neurofilament, and neuron-specific enolase (NSE), to further validate whether H80 cells exhibit fully functional neuronal behavior. H80 cells also require further characterization to determine whether they correspond to distinct neuronal lineages, including dopaminergic, glutamatergic, GABAergic, or cholinergic neurons. Moreover, it remains unclear whether neuronal protein expression in H80 cells arises from intrinsic differentiation potential, genetic reprogramming, or adaptive responses to the tumor microenvironment. Elucidating these mechanisms will be essential for defining the broader biological significance of our observations. However, despite limitations, our study provides compelling evidence that H80 cells possess neuronal lineage features, as demonstrated by the expression of NeuN, MAP2, and Tau. These findings expand the characterization of H80 cells and underscore their potential as a hybrid model system at the intersection of glioma biology and neurodegenerative research. Nevertheless, some of the salient findings have been confirmed in well-established neuronal models, such as SH-SY5Y neuronal cell line, and 3D models, such as spheroid and organoids containing either neuronal cell line (SHSY5Y) or iPSCs-derived neurons, respectively, which substantially enhance the rigor and robustness of the findings.

## Conclusion

These findings collectively support a unified model in which RSK1 functions as a central signaling hub integrating diverse upstream perturbations into a common downstream outcome, pathological Tau phosphorylation. The convergence of HIV- and cocaine-induced signaling on RSK1 provides a mechanistic framework for understanding how viral infection and substance abuse jointly exacerbate neurodegenerative processes, particularly in the context of neuroHIV. Importantly, this RSK1-driven mechanism appears to operate independently of GSK3β, indicating the presence of an additional regulatory layer beyond the established role of GSK3β in Tau phosphorylation. In the context of cocaine exposure, this pathway is further reinforced through coordinated activation of AKT, supporting a dual-axis signaling network that integrates survival and stress-responsive pathways into a shared pathological outcome. This study provides direct evidence that RSK1 is not merely associated with but is functionally required for Tau phosphorylation in the setting of viral and substance-induced neuronal stress. By identifying RSK1 as a dominant and convergent driver of Tau pathology, our work redefines the regulatory hierarchy of Tau-directed kinase signaling and uncovers a critical, previously underappreciated mechanism underlying neurodegeneration. From a translational perspective, the identification of RSK1 as a dominant and druggable driver of Tau pathology has important implications. The ability of RSK1 inhibition to reverse Tau hyperphosphorylation and restore kinase balance suggests that targeting this pathway may offer a viable therapeutic strategy for mitigating Tauopathy in HIV-associated neurocognitive disorders, as well as in broader neurodegenerative conditions. More broadly, these results provide mechanistic clarity to longstanding clinical observations linking HIV infection and stimulant use with accelerated cognitive decline, and position RSK1 as a promising point of intervention for preventing or slowing neurodegeneration. Future studies will focus on evaluating whether pharmacologic inhibition of RSK1 in physiologically relevant preclinical models, including humanized mouse systems, can attenuate neuroinflammation, suppress Tau pathology, and preserve neuronal function. Such investigations will be critical for establishing the therapeutic viability of RSK1-targeted interventions and may ultimately pave the way for novel treatment strategies aimed at preventing or slowing neurodegeneration in HAND and related Tau-associated disorders.

## Acknowledgement

We thank the AIDS Research and Reagent Program, Division of AIDS, National Institute of Allergy, and Infectious Diseases, US National Institutes of Health. We thank Dr. Jonathan Karn and his laboratory for providing the C20 human microglial cell line. We acknowledge the NIH HIV Reagent Program (Division of AIDS, NIAID, NIH) and Dr. Douglas Richman for providing MT 4 cells (ARP 120). This study utilized services offered by core facilities of Thomas Jefferson University (FACS and Imaging) and the Comprehensive NeuroHIV Center (CNHC) at Temple University Lewis Katz School of Medicine. Moreover, we would like to thank the Center for Translational Medicine, Thomas Jefferson University, including all staff members for their technical support and assistance in conducting the experiments for this study.

## Authors’ Contribution

Conceptualization, A.L.S. and M.T.; methodology, A.L.S. and I.K.S.; software, A.L.S., I.K.S., and M.T.; validation, A.L.S., I.K.S., and M.T.; formal analysis, A.L.S., I.K.S., U.P.N., and M.T.; investigation, A.L.S., I.K.S., U.P.N., and M.T.; data curation, A.L.S., I.K.S., U.P.N., and M.T.; writing—original draft preparation and review, A.L.S., I.K.S., U.P.N., and M.T.; project supervision and funding acquisition, M.T.; all authors have read and approved the final version of the manuscript.

## Ethics statement

No ethical approval required

## Generative AI statement

The authors declare that no generative AI was used in the creation of this manuscript

## Resource availability

### Lead contact

Requests for further information and resources should be directed to and will be fulfilled by the lead contact, Mudit Tyagi.

### Materials availability

This study did not generate new unique reagents.

### Data and code availability

- This paper does not report original code.
- Any additional information required to reanalyze the data reported in this paper is available from the lead contact upon request.

## Key resources table

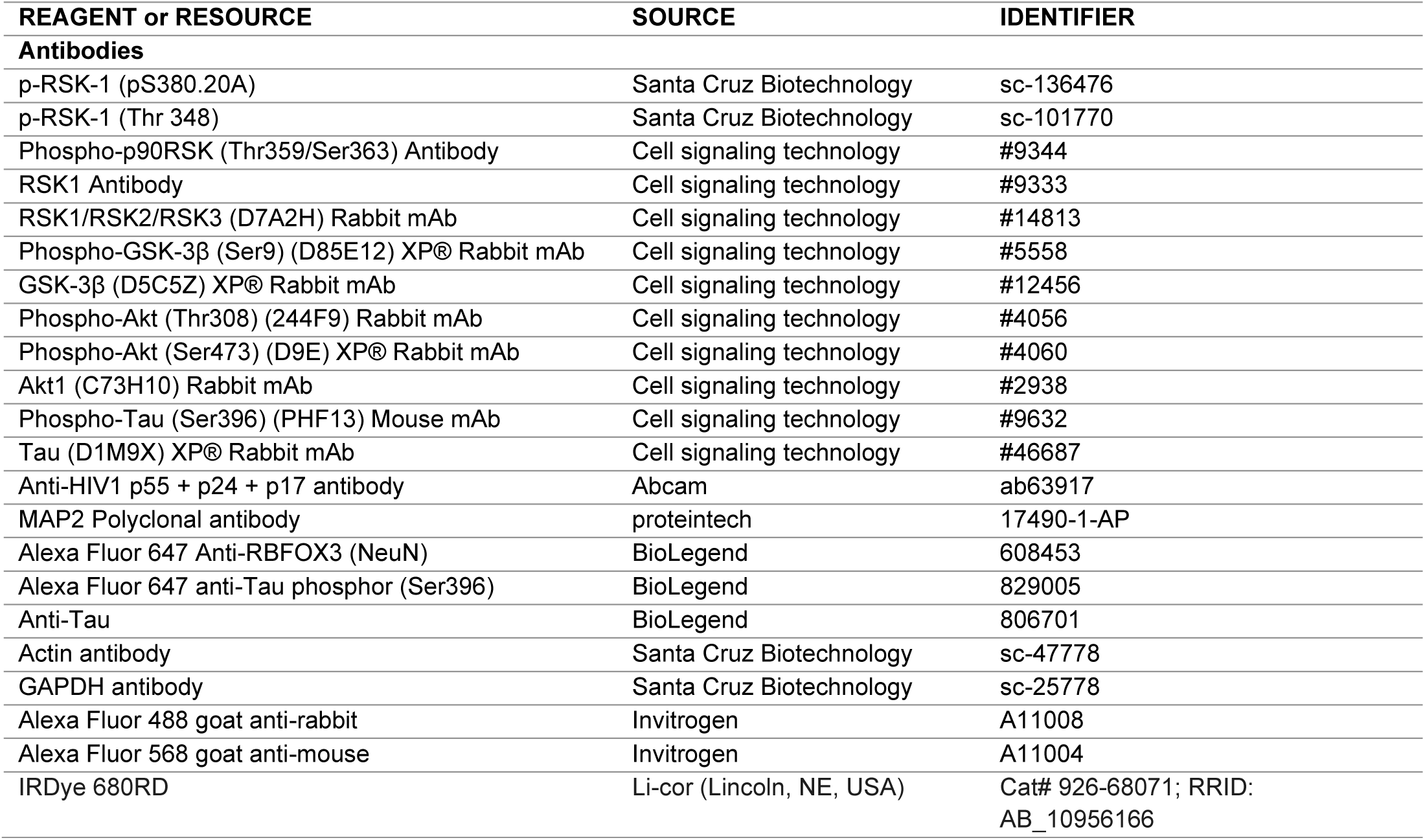
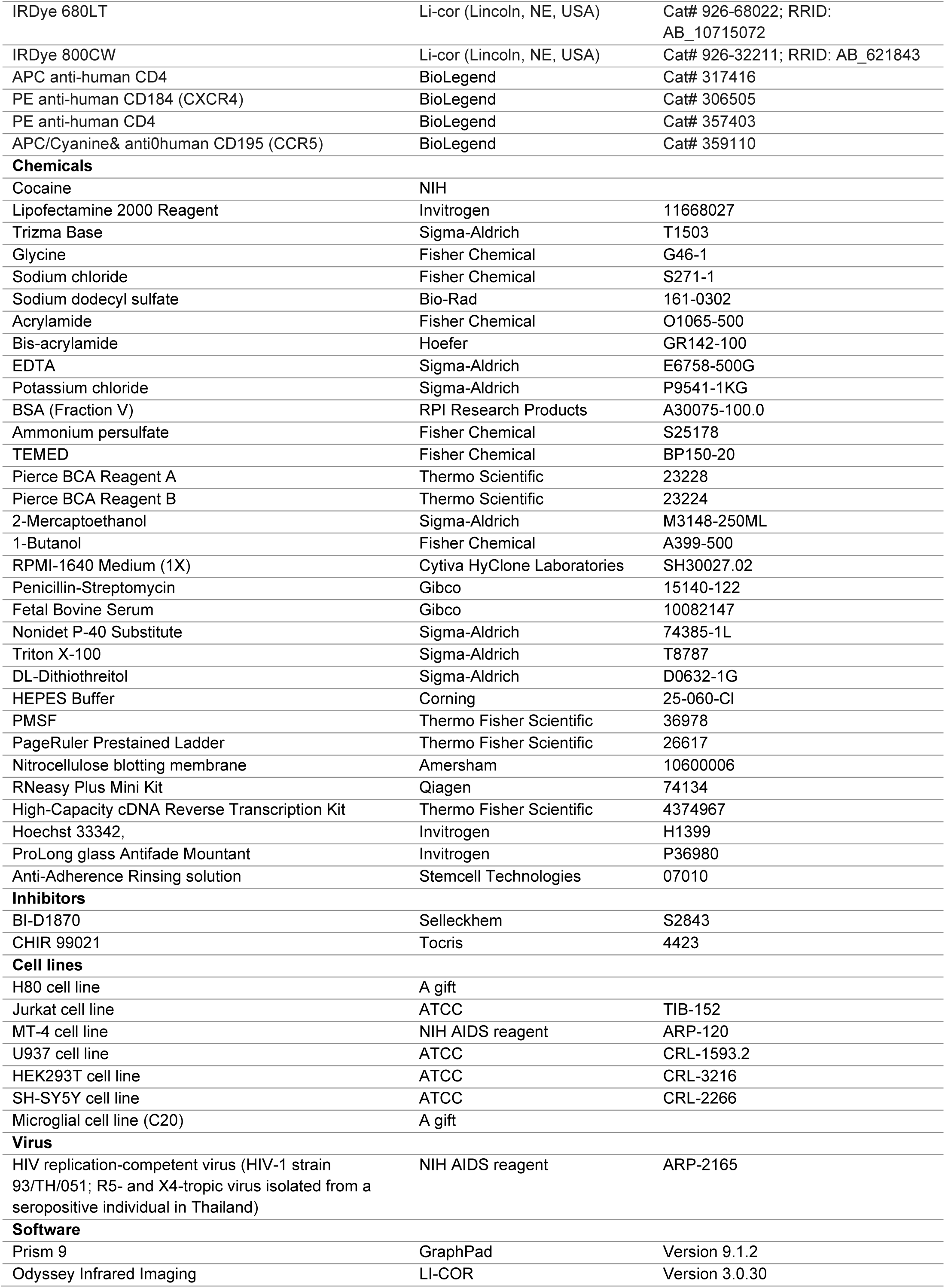
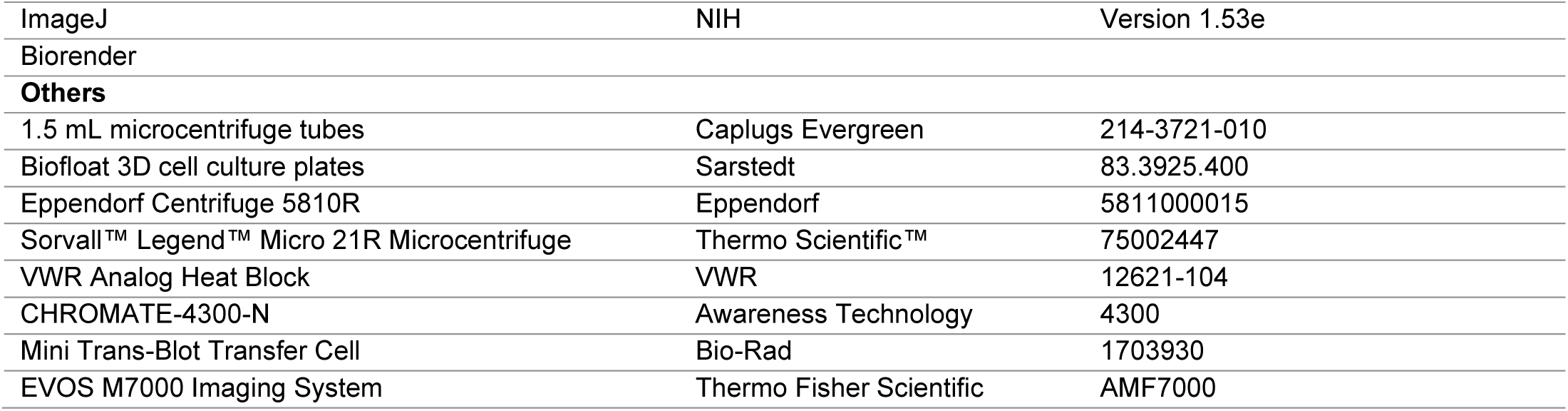

| REAGENT or RESOURCE | SOURCE | IDENTIFIER |
| --- | --- | --- |
| **Antibodies** |  |  |
| p-RSK-1 (pS380.20A) | Santa Cruz Biotechnology | sc-136476 |
| p-RSK-1 (Thr 348) | Santa Cruz Biotechnology | sc-101770 |
| Phospho-p90RSK (Thr359/Ser363) Antibody | Cell signaling technology | #9344 |
| RSK1 Antibody | Cell signaling technology | #9333 |
| RSK1/RSK2/RSK3 (D7A2H) Rabbit mAb | Cell signaling technology | #14813 |
| Phospho-GSK-3β (Ser9) (D85E12) XP® Rabbit mAb | Cell signaling technology | #5558 |
| GSK-3β (D5C5Z) XP® Rabbit mAb | Cell signaling technology | #12456 |
| Phospho-Akt (Thr308) (244F9) Rabbit mAb | Cell signaling technology | #4056 |
| Phospho-Akt (Ser473) (D9E) XP® Rabbit mAb | Cell signaling technology | #4060 |
| Akt1 (C73H10) Rabbit mAb | Cell signaling technology | #2938 |
| Phospho-Tau (Ser396) (PHF13) Mouse mAb | Cell signaling technology | #9632 |
| Tau (D1M9X) XP® Rabbit mAb | Cell signaling technology | #46687 |
| Anti-HIV1 p55 + p24 + p17 antibody | Abcam | ab63917 |
| MAP2 Polyclonal antibody | proteintech | 17490-1-AP |
| Alexa Fluor 647 Anti-RBFOX3 (NeuN) | BioLegend | 608453 |
| Alexa Fluor 647 anti-Tau phosphor (Ser396) | BioLegend | 829005 |
| Anti-Tau | BioLegend | 806701 |
| Actin antibody | Santa Cruz Biotechnology | sc-47778 |
| GAPDH antibody | Santa Cruz Biotechnology | sc-25778 |
| Alexa Fluor 488 goat anti-rabbit | Invitrogen | A11008 |
| Alexa Fluor 568 goat anti-mouse | Invitrogen | A11004 |
| IRDye 680RD | Li-cor (Lincoln, NE, USA) | Cat# 926-68071; RRID:<br>AB_10956166 |
| IRDye 680LT | Li-cor (Lincoln, NE, USA) | Cat# 926-68022; RRID: AB_10715072 |
| IRDye 800CW | Li-cor (Lincoln, NE, USA) | Cat# 926-32211; RRID: AB_621843 |
| APC anti-human CD4 | BioLegend | Cat# 317416 |
| PE anti-human CD184 (CXCR4) | BioLegend | Cat# 306505 |
| PE anti-human CD4 | BioLegend | Cat# 357403 |
| APC/Cyanine& anti0human CD195 (CCR5) | BioLegend | Cat# 359110 |
| **Chemicals** |  |  |
| Cocaine | NIH |  |
| Lipofectamine 2000 Reagent | Invitrogen | 11668027 |
| Trizma Base | Sigma-Aldrich | T1503 |
| Glycine | Fisher Chemical | G46-1 |
| Sodium chloride | Fisher Chemical | S271-1 |
| Sodium dodecyl sulfate | Bio-Rad | 161-0302 |
| Acrylamide | Fisher Chemical | O1065-500 |
| Bis-acrylamide | Hoefer | GR142-100 |
| EDTA | Sigma-Aldrich | E6758-500G |
| Potassium chloride | Sigma-Aldrich | P9541-1KG |
| BSA (Fraction V) | RPI Research Products | A30075-100.0 |
| Ammonium persulfate | Fisher Chemical | S25178 |
| TEMED | Fisher Chemical | BP150-20 |
| Pierce BCA Reagent A | Thermo Scientific | 23228 |
| Pierce BCA Reagent B | Thermo Scientific | 23224 |
| 2-Mercaptoethanol | Sigma-Aldrich | M3148-250ML |
| 1-Butanol | Fisher Chemical | A399-500 |
| RPMI-1640 Medium (1X) | Cytiva HyClone Laboratories | SH30027.02 |
| Penicillin-Streptomycin | Gibco | 15140-122 |
| Fetal Bovine Serum | Gibco | 10082147 |
| Nonidet P-40 Substitute | Sigma-Aldrich | 74385-1L |
| Triton X-100 | Sigma-Aldrich | T8787 |
| DL-Dithiothreitol | Sigma-Aldrich | D0632-1G |
| HEPES Buffer | Corning | 25-060-CI |
| PMSF | Thermo Fisher Scientific | 36978 |
| PageRuler Prestained Ladder | Thermo Fisher Scientific | 26617 |
| Nitrocellulose blotting membrane | Amersham | 10600006 |
| RNeasy Plus Mini Kit | Qiagen | 74134 |
| High-Capacity cDNA Reverse Transcription Kit | Thermo Fisher Scientific | 4374967 |
| Hoechst 33342, | Invitrogen | H1399 |
| ProLong glass Antifade Mountant | Invitrogen | P36980 |
| Anti-Adherence Rinsing solution | Stemcell Technologies | 07010 |
| **Inhibitors** |  |  |
| BI-D1870 | Selleckhem | S2843 |
| CHIR 99021 | Tocris | 4423 |
| **Cell lines** |  |  |
| H80 cell line | A gift |  |
| Jurkat cell line | ATCC | TIB-152 |
| MT-4 cell line | NIH AIDS reagent | ARP-120 |
| U937 cell line | ATCC | CRL-1593.2 |
| HEK293T cell line | ATCC | CRL-3216 |
| SH-SY5Y cell line | ATCC | CRL-2266 |
| Microglial cell line (C20) | A gift |  |
| **Virus** |  |  |
| HIV replication-competent virus (HIV-1 strain 93/TH/051; R5- and X4-tropic virus isolated from a seropositive individual in Thailand) | NIH AIDS reagent | ARP-2165 |
| **Software** |  |  |
| Prism 9 | GraphPad | Version 9.1.2 |
| Odyssey Infrared Imaging | LI-COR | Version 3.0.30 |
| ImageJ | NIH | Version 1.53e |
| Biorender |  |  |
| **Others** |  |  |
| 1.5 mL microcentrifuge tubes | Caplugs Evergreen | 214-3721-010 |
| Biofloat 3D cell culture plates | Sarstedt | 83.3925.400 |
| Eppendorf Centrifuge 5810R | Eppendorf | 5811000015 |
| Sorvall™ Legend™ Micro 21R Microcentrifuge | Thermo Scientific™ | 75002447 |
| VWR Analog Heat Block | VWR | 12621-104 |
| CHROMATE-4300-N | Awareness Technology | 4300 |
| Mini Trans-Blot Transfer Cell | Bio-Rad | 1703930 |
| EVOS M7000 Imaging System | Thermo Fisher Scientific | AMF7000 |

