## Supplemental file for "HIV and Cocaine exposure promote Tau phosphorylation through RSK-1 in a GSK3β-independent manner"

Original Research article

**Figure S1**

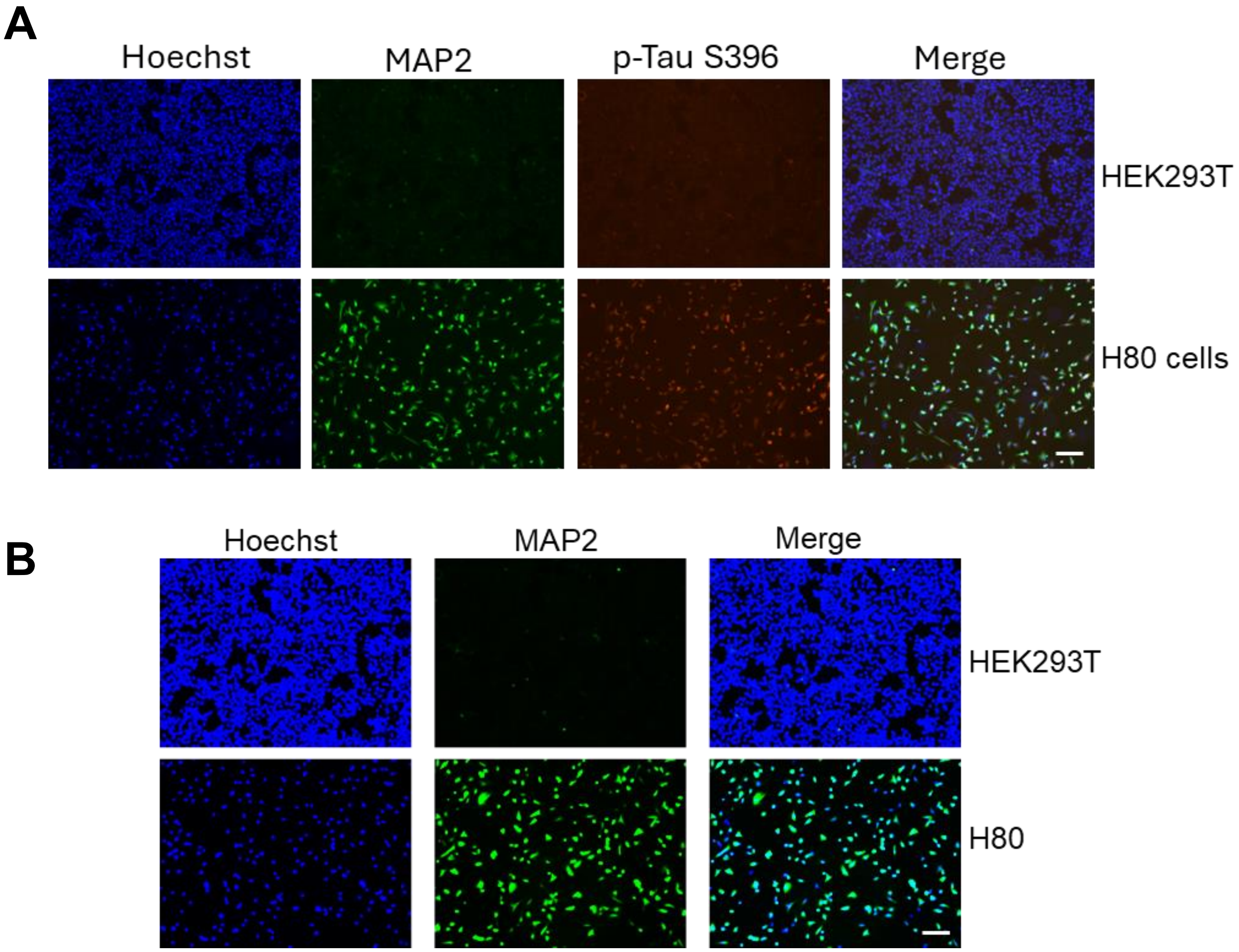


**Figure S1: MAP2 and Tau expression confirm the neuronal identity of H80 cells. (A)** Immunofluorescence staining of H80 cells and HEK293T cells comparing the expression of MAP2 and phosphorylated Tau (p Tau Ser396). MAP2 and p-Tau Ser396 were readily detected in H80 cells but were not detectable in HEK293T cells, indicating neuronal specificity of these markers. These data collectively support that H80 cells display molecular and structural features characteristic of the neuronal lineage. **(B)** Immunofluorescence reanalysis of microtubule associated protein 2 (MAP2) in H80 cells and HEK293T cells (non neuronal control). H80 cells exhibited a pronounced filamentous MAP2 signal consistent with dendritic architecture, whereas HEK293T cells showed no detectable MAP2 staining under identical conditions. Scale bar: 10 μm. **Related to Fig. 1.**

**Figure S2**

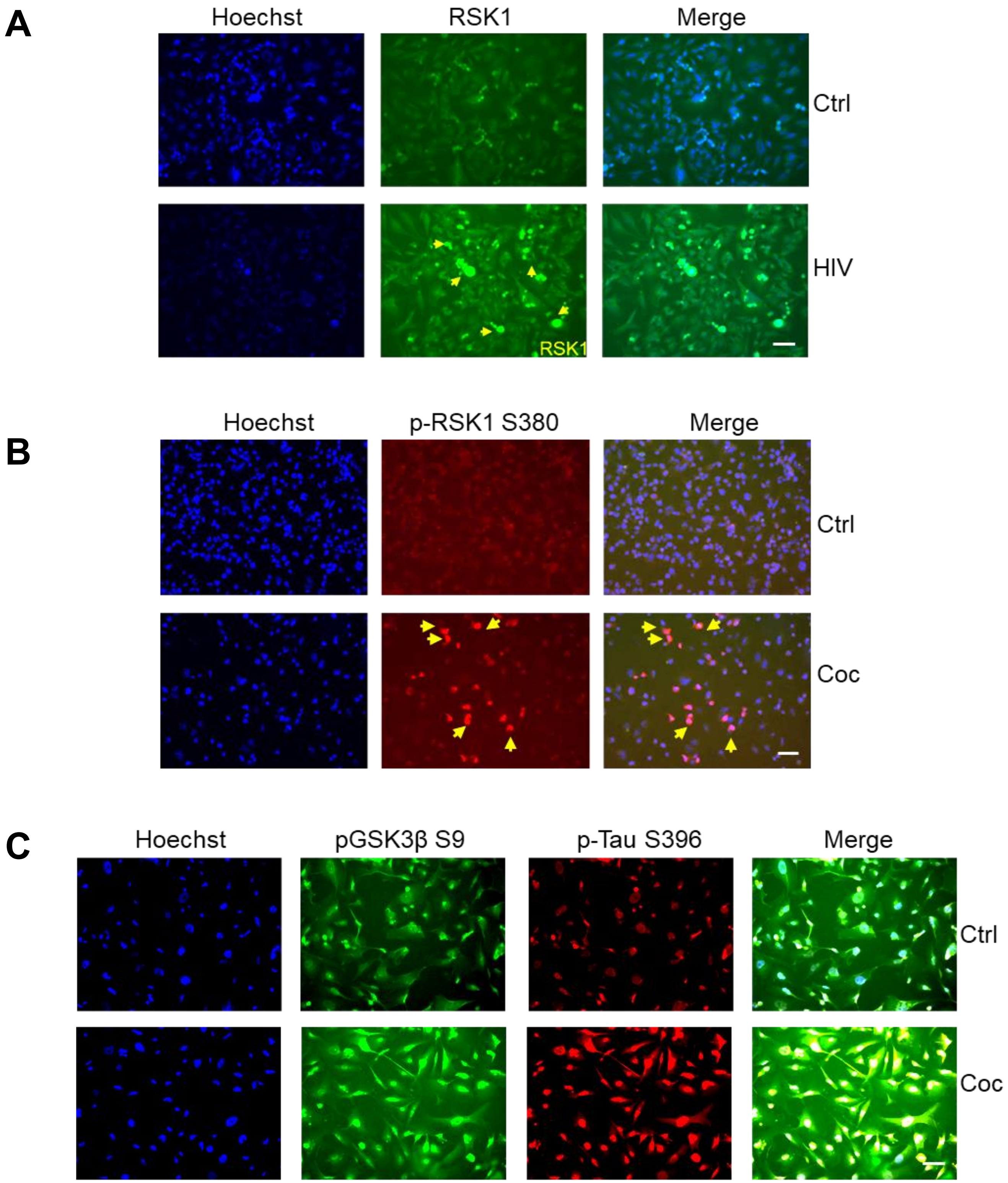


**Figure S2: Chronic HIV and cocaine exposure upregulates and activates RSK1, inactivates GSK3β, and promotes Tau phosphorylation in H80 cells.** (**A**) Immunofluorescence analysis of H80 cells chronically exposed to HIV revealed a marked increase in RSK1 immunoreactivity compared with untreated controls, indicating HIV induced upregulation of RSK1 expression. (**B**) Immunofluorescence analysis of H80 cells exposed to cocaine demonstrated a robust increase in p-RSK1 Ser380, consistent with cocaine induced activation of RSK1. (**C**) Cocaine treatment also elevated phosphorylation of GSK3β at Ser9, indicative of functional GSK3β inactivation, and concurrently increased Tau phosphorylation. **Related to Figures 3 and 4.**

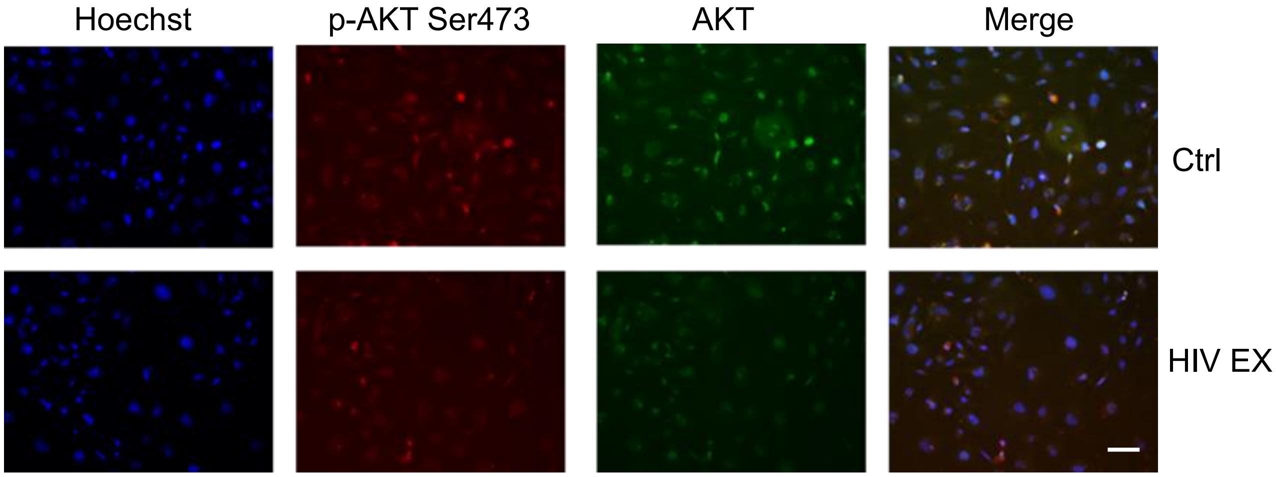


**Figure S3. HIV exposure does not activate AKT signaling.** Immunofluorescence analysis of H80 cells exposed to HIV revealed no detectable change in AKT phosphorylation at Ser473 (p-AKT-Ser473) compared to control, indicating that HIV does not activate the AKT pathway. Nuclear staining was performed using Hoechst. Total AKT levels remained unchanged, confirming that HIV exposure alone does not modulate AKT activation. Scale bar: 10 µm. **Related to main Figure. 5**.

**Figure S4**

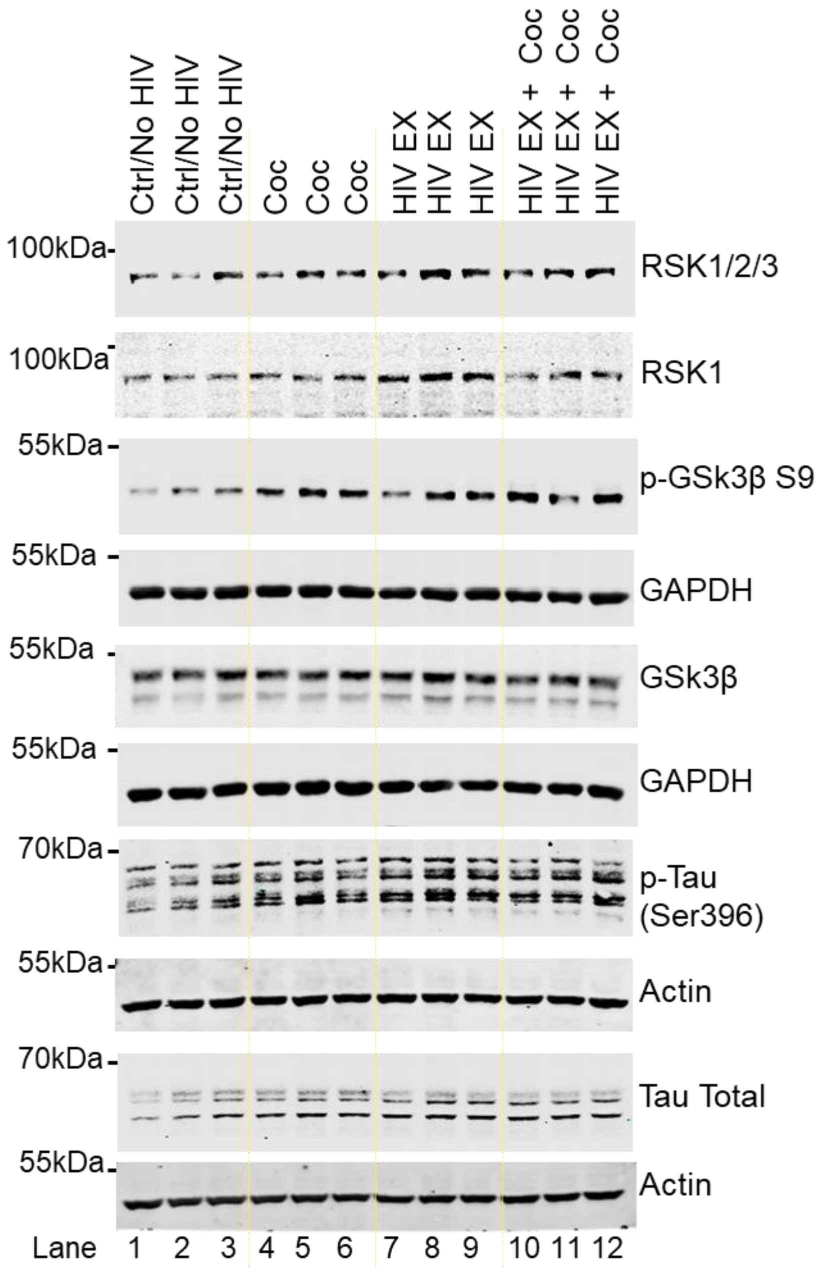


**Figure S4: Grayscale version of Fig. 6. Cocaine and HIV exposure increase Tau phosphorylation independent of GSK3β activation.** Immunoblot analysis of H80 cells exposed for 48 h to cocaine, HIV virions, or their combination revealed a marked increase in Tau phosphorylation at Ser396 (p-Tau Ser396), a pathological Tau epitope, compared with untreated control cells. Parallel analysis showed increased phosphorylation of GSK3β at Ser9 (p-GSK3β Ser9) under all treatment conditions, consistent with functional inhibition of GSK3β. Total GSK3β and total Tau protein levels remained unchanged across conditions. This panel represents a grayscale rendering of the corresponding color immunoblot shown in main **Figure. 6**.

**Figure S5**

**A**

**B**

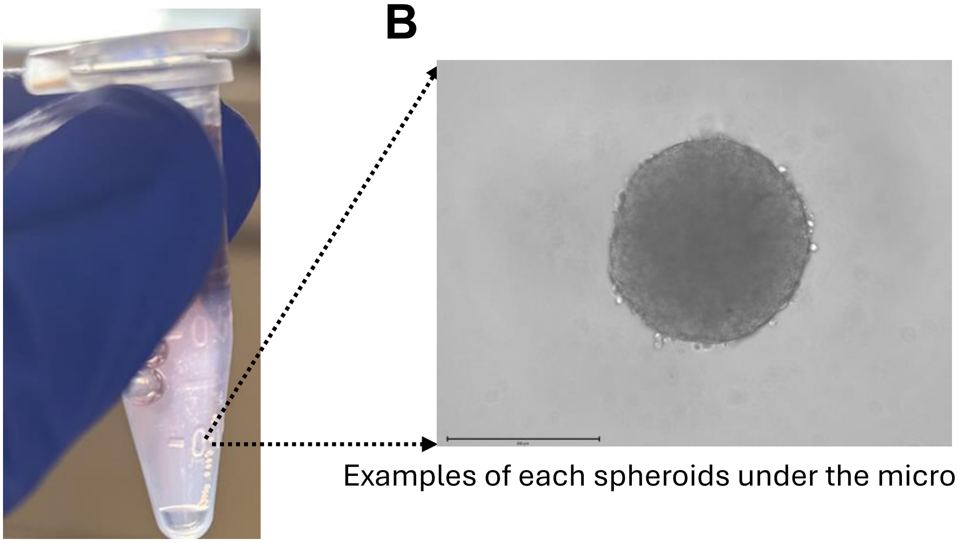

Examples of each spheroids under the microscope

Examples of spheroids in 1.5 ml microcentrifuge tube

**Figure S5. Spheroid Experimental Design** For each experimental condition, 96 spheroids were generated from 96 individual wells, with each spheroid treated independently. Spheroids from each condition were then pooled into a single tube (n = 24 spheroids per condition X 4= 96). The experimental groups were as follows: Control (Ctrl): 24 untreated spheroids; Cocaine: 24 spheroids individually treated with cocaine; HIV: 24 spheroids individually exposed to HIV; HIV + Cocaine: 24 spheroids individually exposed to both HIV and cocaine. **(A)** Representative image of 24 pooled spheroids collected in a 1.5-mL microcentrifuge tube. **(B)** Microscopy image showing individual spheroids prior to pooling. Scale bar : 500 µm **Related to main Figure 9**.

**C**

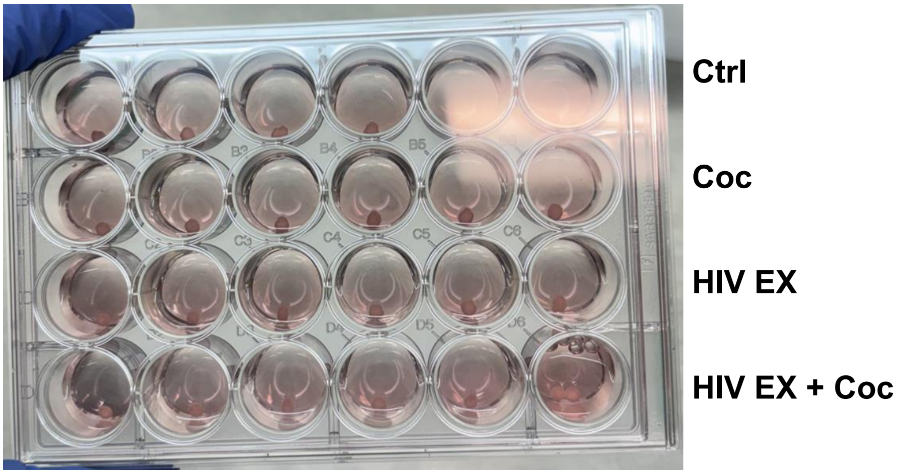


**Figure S5. (C) Organoid culture under four experimental conditions.** Human brain organoids were maintained in 24-well plate cultures and subjected to four treatment conditions: Ctrl, Cocaine, HIV, and HIV + Cocaine. Each condition represents an independently treated organoid culture. **Related to main Figure 9.**
